# Protein overabundance is driven by growth robustness

**DOI:** 10.1101/2024.08.14.607847

**Authors:** H. James Choi, Teresa W. Lo, Kevin J. Cutler, Dean Huang, W. Ryan Will, Paul A. Wiggins

**Affiliations:** Department of Physics, University of Washington, Seattle, Washington 98195, USA; Department of Laboratory Medicine and Pathology, University of Washington, Seattle, Washington 98195, USA; Department of Microbiology, University of Washington, Seattle, Washington 98195, USA; Department of Bioengineering, University of Washington, Seattle, Washington 98195, USA

## Abstract

Protein expression levels optimize cell fitness: Too low an expression level of essential proteins will slow growth by compromising essential processes; whereas overexpression slows growth by increasing the metabolic load. This trade-off naïvely predicts that cells maximize their fitness by sufficiency, expressing just enough of each essential protein for function. We test this prediction in the naturally-competent bacterium *Acinetobacter baylyi* by characterizing the proliferation dynamics of essential-gene knockouts at a single-cell scale (by imaging) as well as at a genome-wide scale. In these experiments, cells proliferate for multiple generations as target protein levels are diluted from their endogenous levels. This approach facilitates a proteome-scale analysis of the fitness landscape with respect to protein abundance. We find that most essential proteins are subject to a threshold-like fitness landscape: growth is independent of protein abundance above a critical threshold and arrests below that threshold. We have recently analyzed the implications of this landscape for growth robustness. Confirming signature predictions of this model, we find that (i) roughly 70% of essential proteins are overabundant, (ii) overabundance increases as the expression level decreases and (iii) the lowest abundance proteins are in vast excess (*>*10×) of what is required for growth in the typical cell. These results reveal that robustness plays a fundamental role in determining the expression levels of essential genes and that overabundance is a key mechanism for ensuring robust growth.

---

Understanding the rationale for protein expression levels is a fundamental question in biology with broad implications for understanding cellular function [1]. Measured expression levels appear to be paradoxically both *optimal* and *overabundant*. For instance, repeated investigations support the idea that gene expression levels optimize cell fitness [2, 3]. Since the overall metabolic cost of protein expression is large [4, 5], fitness optimization would seem to imply that protein levels should satisfy a *Goldilocks condition*: Expression levels should be *just high enough* to achieve the required protein activity [6, 7]. However, a range of approaches suggest that many essential genes are expressed in vast excess of the levels required for function [7–9]. How can expression levels be at once optimal with respect to fitness as well as in excess of what is required for function?

The cell faces a complex regulatory challenge: Even in a bacterium, there are between four and six hundred essential proteins, each of which is required for growth [10]. How does the cell ensure the robust expression of each essential factor? We recently argued that the stochasticity of gene expression processes fundamentally shape the principles of central dogma regulation, including the optimality of protein overabundance [11]. Specifically, we proposed a quantitative model, the Robustness-Load Trade-Off (RLTO) model, which makes a parameter-free prediction of protein overabundance as a function of gene transcription level [11]. The optimality of overabundance can be understood as the result of a highly-asymmetric fitness landscape: the fitness cost of essential protein underabundance, which causes the arrest of essential processes, is far greater than the fitness cost of essential protein overabundance, which leads to slow growth by increasing the metabolic load. However, critical model assumptions and predictions remain untested which is the motivation for the current study. Here, we will quantitatively measure the fitness landscape with repect to protein abundance and determine the level of overabundance for all essential proteins in the bacterium *Acinetobacter baylyi*.

## RESULTS

### Natural competence facilitates knockout-depletion

To characterize the fitness landscape for essential gene expression, we must deplete the levels of essential proteins. Both degron- and CRISPRi-based approaches have been applied; however, these approaches require careful characterization of protein levels [8, 12–15] and could introduce significant cell-to-cell variation at the single-cell scale [16], on top of the endogenous gene expression noise, which further obscures the underlying fitness landscape. To circumvent these difficulties, we will use an alternative approach: *knockout-depletion* in the naturally competent bacterium *A. baylyi* ADP1 [17, 18]. In this approach, cells are transformed with a *geneX∷kan* knockout cassettes targeting essential gene X, carrying a kanamycin resistance allele Km^*R*^. (See Fig. 1A.) Cells that are not transformed arrest immediately on selective media. The crux of the approach is that transformants remain transiently *geneX*^+^, due to the presence of already synthesized target protein X, even after the transcription of the target *geneX* stops. Growth can continue, diluting protein X abundance, as long as this residual abundance remain sufficient for function. The success of the knockout-depletion approach is dependent on the extremely high transformation efficiency of *A. baylyi*.

**FIG. 1.**
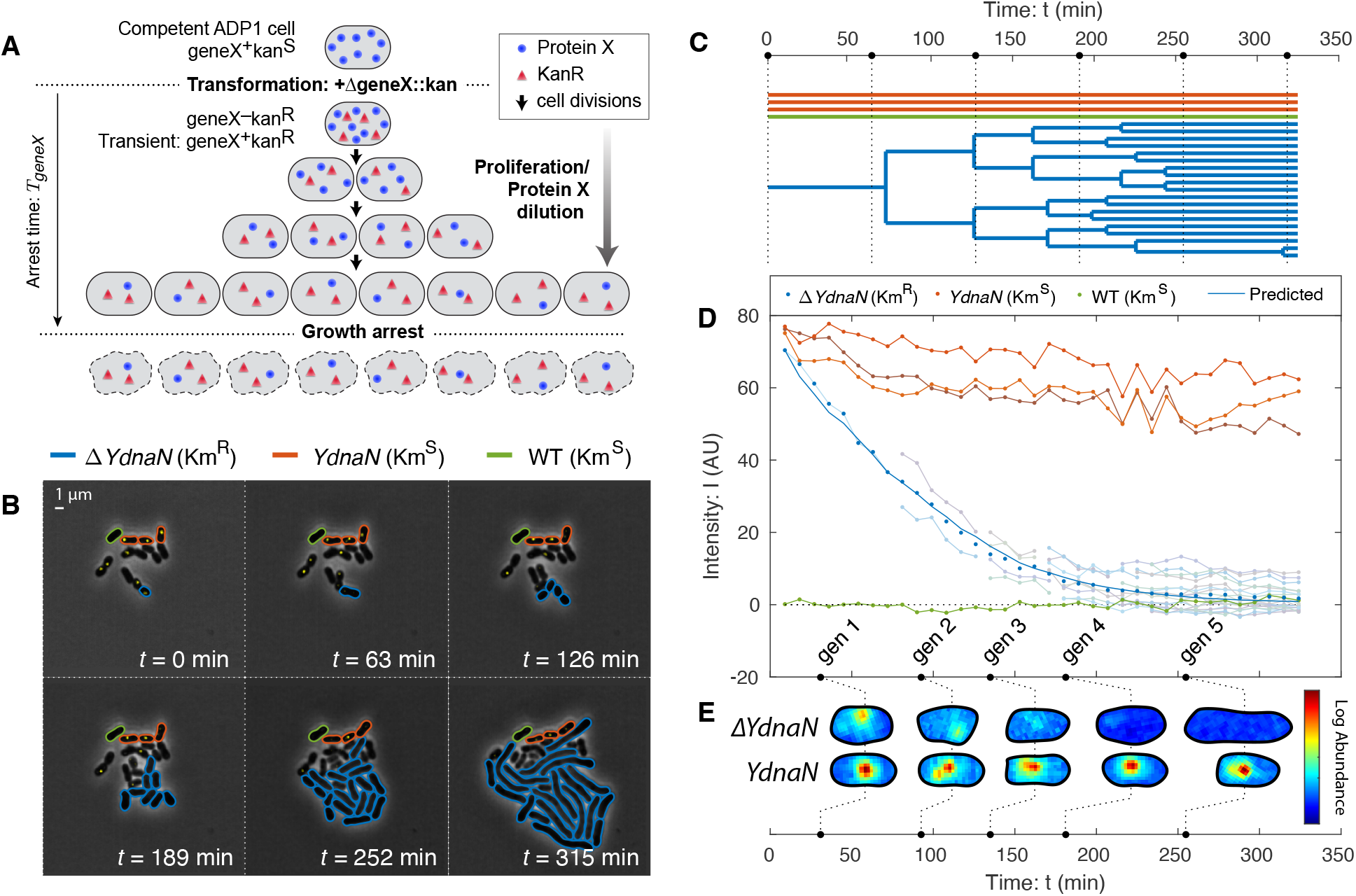
Knockout-depletion experiments. Panel A: Experimental schematic. Competent ADP1 cells are transformed with Δ*geneX∷kan*. Untransformed cells arrest immediately on selective media. Transformed cells proliferate, but cease protein X expression (blue circles) while expressing Kan (red triangles). Existing protein X abundance is diluted as cells proliferate. For essential genes, cell growth continues until protein levels are diluted to the threshold level required for growth, after which growth arrests. **Panel B & C: Visualization of knockout depletion**. The fluorescent fusion *YPet-dnaN* to essential gene *dnaN* is knocked out at *t* = 0. Cell proliferation is visualized using phase-contrast microscopy while protein abundance is measured by fluorescence microscopy (yellow). Transformed cells (Δ*YdnaN*, blue) have a Km^*R*^ allele and can proliferate over several generations before arrest; however, untransformed cells (*YdnaN*, orange) and wild-type cells (WT, green) were both kanamycin sensitive and therefore arrested immediately. (See Supplemental Material Sec. 4 B 2.) **Panel C: Lineage tree**. Black dotted lines represent time points shown in Panel B. **Panel D: Target protein is diluted by proliferation**. Protein concentration is measured by integrated fluorescence. Arrested *YdnaN* cells maintain protein abundance, whereas proliferating transformed cells (Δ*YdnaN*, blue) show growth-induced protein depletion. The protein concentration over all transformed progeny (blue points) are consistent with the dilution-model prediction (solid blue). **Panel E: Protein function is robust to dilution**. Representative single-cell images of transformed (Δ*YdnaN*) and untransformed (*YdnaN*) cells are shown for successive time points. The YPet-DnaN fusion shows punctate localization, consistent with function, even as protein abundance is depleted. No puncta are observed in the last generation and the cells form filaments, consistent with the replication arrest phenotype.

### Target proteins are depleted by dilution

A key untested assumption in the experimental design of the knockout-depletion approach is that target protein translation stops after transformation, and that the protein abundance is depleted by dilution. The model predicts that the protein concentration is:

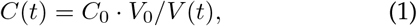

where *C*_0_ and *V*_0_ are the concentration and volume of the progenitor cell at deletion and *V* (*t*) is the total volume of the progeny. To test the predicted protein depletion hypothesis, we designed a knockout-depletion experiment to target a protein we had previously studied that can be visualized using a fluorescent fusion and whose localization is activity dependent: the essential replication gene *dnaN*, whose gene product is the *β* sliding clamp [19–21]. We constructed a N-terminal fluorescent fusion to *dnaN* using YPet in *A. baylyi* at the endogenous locus. The resulting mutant (YdnaN) had no measurable growth defect under our experimental conditions. We then knocked out the *YPet-dnaN* fusion, yielding Δ*dnaN*, and characterized the protein levels by quantifying YPet-DnaN abundance by fluorescence.

The experimental design of knockout-depletion assays (Fig. 1A) can fail by a number of distinct mechanisms: (i) Transient growth of kanamycin-sensitive cells on selective media could be misinterpreted as transient growth of transformants. Three lines of evidence argue against this possibility. No transient growth is observed in cells that are not transformed with fragments without the *kan* gene. Furthermore, progeny of cells transformed with *kan*-carrying knockout cassettes exhibit consistent phenotypes (*e*.*g*. the cell lysis phenotype for Δ*murA*). Finally, Kan expression appears to be overabundant itself, as discussed in Supplementary Material Sec. 1 A.

(ii) Competent cells could exhibit protein expression in vast excess of log-phase cells. To rule out this possibilit y, we measured YPet-DnaN abundance in competent and wild-type cells. We observed that competent cells in out growth have comparable protein abundance to logphase cells. (See Supplementary Material Sec. 1 B.) (iii) Another important consideration is the rapidity of transformation and the recombination processes. If the DNA target sequence is not rapidly degraded and expression continues, transient growth could be the consequence of this continuing expression. To rule out this possibility, we determined the dilution rate in four micro-colonies and in each case it was consistent with the rate cell elongation. (We note that two roughly-canceling competing effects limit the precision of this analysis: photobleaching and the fluorescence from neighboring wildtype cells. See Supplementary Material Sec. 1 C.) In additional support to experimental design, dilution is observed immediately on the first cell division, suggesting that the target gene is either rapidly degraded or at least cannot be transcribed. (See Fig. 1D.)

(iv) Another complicating scenario is the possibility that the protein is not partitioned equally between daughter cells. Contrary to this possibility, we have previously studied the partitioning of nearly all localized protein in *Escherichia coli* and quantified protein partitioning between daughter cells. Only proteins localized in polar puncta were asymmetrically partitioned [22]. Consistent with these measurements in *E. coli, A. baylyi* partitions Ypet-DnaN nearly equally between daughters early in proliferation. Late in proliferation a technical limitation, fluorescence from neighboring wild-type cells, limits the ability to quantitatively measure partitioning; however, growth arrest in progeny after the initial cell division appears synchronous, consistent with equipartition of mother-cell protein. (See Fig. 1D.)

Observations (i)-(iv) confirm that proteins are diluted consistent with the experimental design shown schematically in Fig. 1A.

### Replication persists during DnaN depletion

A key subhypothesis of the overabundance model for transient growth is that target protein function continues as the target protein abundance is depleted. An alternative hypothesis for transient growth of the Δ*dnaN* strain is a high initial chromosomal copy-number that is partitioned between daughter cells, even after the replication process itself arrests due to target protein depletion [4, 23]. The imaging-based knockout-depletion experiment tests this hypothesis as well. The localization of DnaN is dependent on activity: During ongoing replication, DnaN is localized in puncta corresponding to replisomes, whereas in the absence of active replication, DnaN has diffuse localization [19–21, 24, 25]. During the knockout-depletion experiment, we observed YPet-DnaN puncta persist as the targeted fusion was depleted (Fig. 1DE), consistent with replication activity after dilution. Only after the YPet-DnaN puncta disappear do the cells begin to adopt the Δ*dnaN* phenotype: cell filamentation (Fig. 1BE). We therefore conclude that function (replication) is robust to significant target protein (DnaN) dilution.

### Other essential knockouts undergo transient growth

To understand the generic consequences of essential protein depletion, we used the imaging-based knockout-depletion experiments to explore essential genes with a range of functions. We initially targeted four essential genes: the replication initiation regulator gene *dnaA* (Supplementary Material Fig. S4 and Movie S5-6), the beta-clamp gene *dnaN* (Supplementary Material Fig. S5 and Movie S7-8), the cell-wall-synthesis gene *murA* (Supplementary Material Fig. S7 and Movie S9-10), and septation-related gene *ftsN* (Supplementary Material Fig. S6 and Movie S11-12), as well as a nonessential IS element with no phenotype as a negative control (Movie S3-4). In each case, transformants continued to proliferate through multiple cell-cycle durations [17] and are therefore consistent with the essential protein overabundance hypothesis. However, in Ref. [17], we were unable to perform a quantitative single-cell analysis of these time-lapse experiments since existing segmentation packages failed to segment the observed morphologies [26]. We therefore developed the *Omnipose* package, which facilitated quantitative analysis of the growth dynamics with single-cell resolution [26]. (See Figs. 1B and 2A.)

### The fitness landscape is highly asymmetric

A key input to the RLTO model is the fitness landscape (growth rate) as a function of protein abundance. Omnipose segmentation facilitates the measurement of single-cell growth rates from the time-lapse imaging experiments. We focus first on the single-cell areal growth rate:

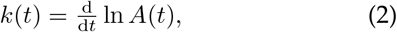

where *A*(*t*) is the area of the cell at time *t*. This areal growth rate is more convenient than a cell-length based rate since we avoid the necessity of defining cell length for unusual cell morphologies like those observed in the Δ*murA* mutant. Fig. 2B shows representative knockout-depletion dynamics of cell area for the essential-gene target *murA*. The log slope remains constant for multiple generations, consistent with a constant growth rate, even as the gene targeted is depleted over multiple cell cycles. By combining the dilution model (Eq. 1) and the growth rate (Eq. 2), a single knockout-depletion measurement determines the growth rate for a range of protein abundances between wild-type abundance and those realized at growth arrest. This fitness landscape is shown for the MurA protein in Fig. 2C. For all four mutants, the areal growth rate is roughly constant for multiple generations before undergoing a rapid transition to growth arrest. (See Supplementary Material Sec. 1.) We emphasize that the predictions of the RLTO model do not depend on a rigorous step-like transition of the growth rate to zero, but rather a rapid decrease in growth rate under a threshold protein abundance as observed for each target [11].

**FIG. 2.**
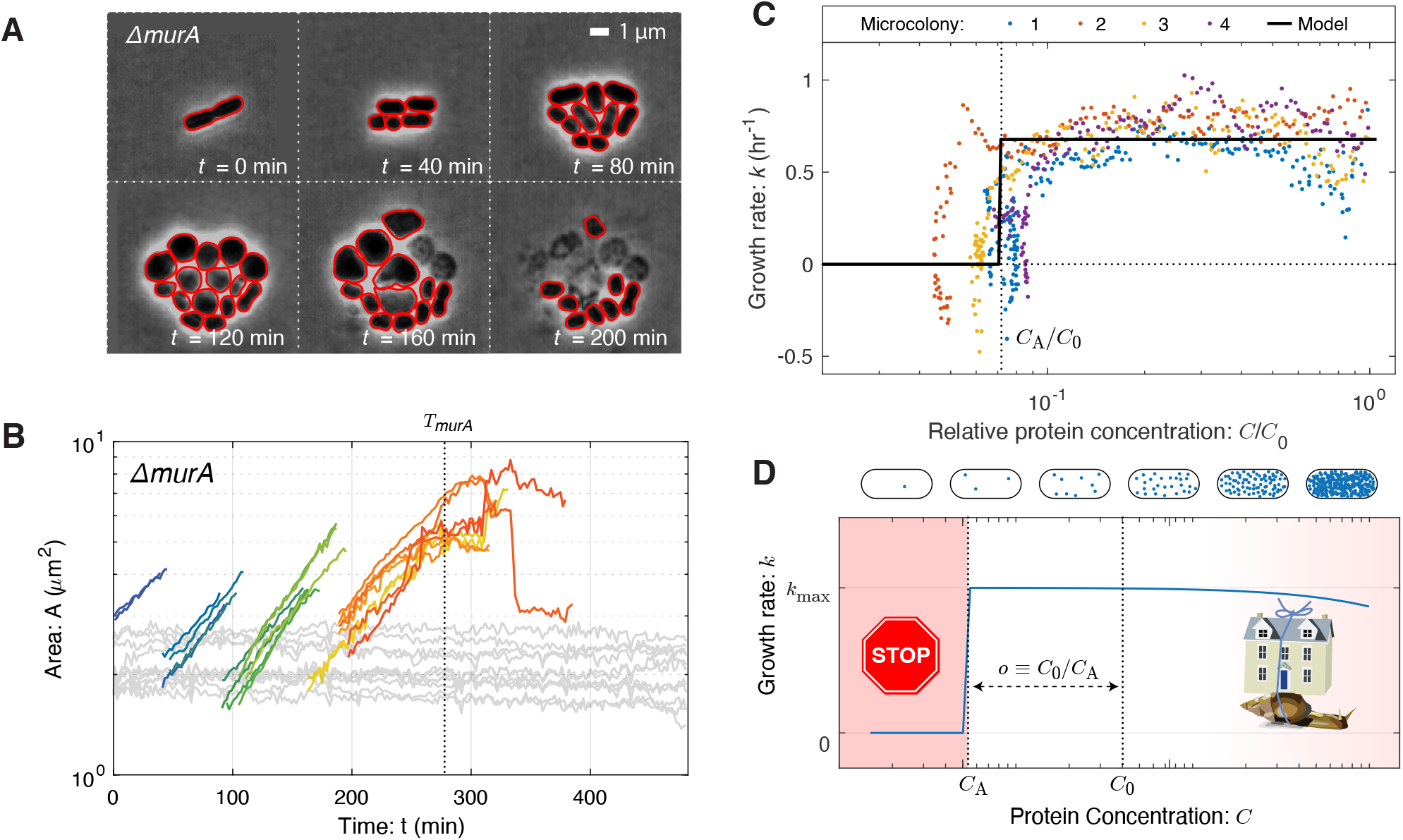
The fitness landscape. Panel A: Visualization of growth in a *murA* knockout. Essential gene *murA* is knocked out at *t* = 0 and cell proliferation is visualized by phase-contrast microscopy. Red outlines represent the Omnipose cell segmentation. Cell proliferation continues for multiple generations after deletion. **Panel B: Quantitative analysis of cell proliferation with single-cell resolution**. Cell area (log scale) as a function of time for the *murA* deletion (colored lines) and representative untransformed cells (light gray). The log-slope represents the single-cell growth rate. The vertical dotted line represents the arrest time at which transformant growth slows to growth arrest. Untransformed cells are arrested throughtout the experiment since the cells are Km^S^. **Panel C: Growth rate as a function of protein depletion for** Δ***murA***. The growth rate is (Eq. 2) is observed to obey a threshold-like-dependence on protein abundance (Eq. 1), transitioning between wild-type growth to arrest at the vertical dotted line. We define the critical dilution as *o* ≡*C*_0_*/C*_*A*_ where *C*_*A*_ is the protein concentration at arrest. Cell-to-cell variation is observed in the arrest of four microcolonies that originate from distinct progenitor cells. The model represents a threshold-like model fit to the growth rate data. **Panel D: The fitness landscape is threshold-like**. Motivated by single-cell growth data, cell fitness is modeled using the Robustness-Load Trade-Off model (RLTO). In the model, there is a metabolic cost of protein expression which favors low expression; however, growth arrests for protein concentration *C* smaller than the threshold level *C*_*A*_ (red). The relative metabolic cost of overabundance is small relative to the cost of growth arrest due to the large number of proteins synthesized, resulting in a highly asymmetric fitness landscape [11].

### Protein overabundance

We will define the overabundance as the ratio of the initial protein concentration (*C*_0_) to the concentration at cell arrest (*C*_*A*_):

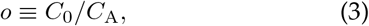

as shown in Fig. 2D. (Supplementary Material Sec. 4 gives a detailed description of the inferred overabundance from single-cell data.) Note that in principle both *C*_0_ and *C*_*A*_ could be determined independently; however, what is measured in the knockout-depletion experiment is the ratio *o*, defined as the overabundance, only.

**TABLE 1.**
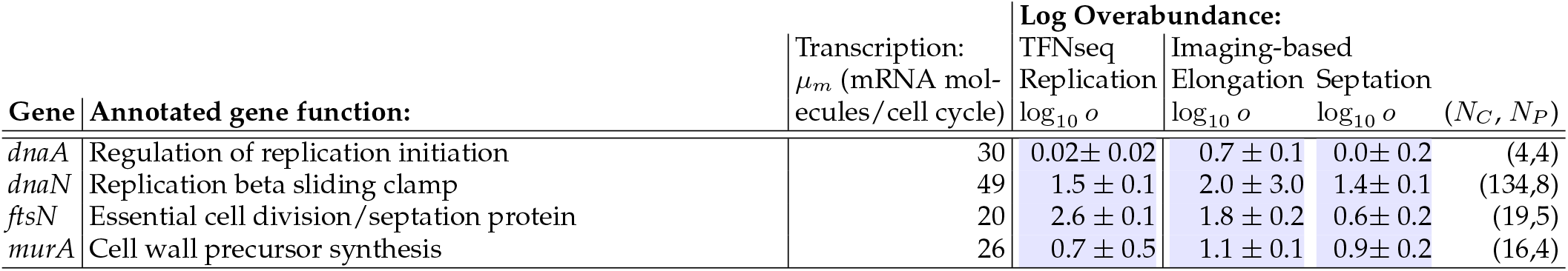
Measured overabundance for sequencing- versus imaging-based approaches. The overabundance was determined by both sequencing- and imaging-based approaches. Each of the three different overabundance measurements is inferred from the arrest of a distinct cellular process: the sequencing-based measurement is determined from dynamics of genomic copy number which depends on replication. For the imaging-based approach, we show two measurements based on different metrics for arrest: The first is based on the arrest of cell elongation, as defined by Eq. 3, and the second is based on the arrest of the septation process, as visualized by microscopy. *N*_*C*_ and *N*_*P*_ are the number of cells analyzed and the number of progenitor cells analyzed in the imaging-based experiments. For sequencing-based measures, errors are estimated from the analysis of replicate experiments. For imaging-based errors, errors are estimated from the variance between arrest times for distinct progenitor cells. (See Sumplementary Material Tab. S1 for a more detailed description of analysis.)

The measured overabundance for the four mutants imaged by microscopy is summarized in Tab. I, using three distinct metrics for growth. We conclude that for each gene, with the exception of *dnaA*, rapid growth continues after the knockout due to the vast overabundance of the target protein.

### The RLTO model predicts protein overabundance

The RLTO model explicitly analyzes the trade-off between growth robustness to noise and metabolic load and predicts the optimal central-dogma regulatory principles [11]. Critically, the model incorporates the observed threshold-like dependence of growth rate on protein abundance (Fig. 2CD). The model quantitatively predicts protein overabundance with a signature feature: high-expression genes have low protein overabundance (*o*≈1) due to the high metabolic cost of increasing expression and low inherent noise of high expression genes; however, low-expression genes have high overabundance (*o*≫1) due to the low metabolic cost of increasing expression and the high inherent noise of low expression genes. (See Supplemental Material Sec. 7 for a more detailed description of the model.)

### TFNseq determines overabundances genome-wide

To test the signature expression-dependent overabudance prediction of the RLTO model, we now transition to a genomic-scale analysis. The Manoil lab developed a TFNseq-approach to knockout-depletion experiments for targeting all genes simultaneously in *A. baylyi* [18]. In short: A genomic library was prepared and mutagenized using a transposon carrying the Km^R^ allele. The resulting DNA was then transformed into *A. baylyi*. The transformants were propagated on selective liquid media and fractions collected every two hours from which genomic DNA was extracted. The transposons were then mapped using TFNseq to generate the relative abundance trajectory for each mutant [18]. (See Fig. 3AB.) We then analyzed each mutant trajectory statistically using three competing growth models: noeffect, sufficiency, and overabundance, using two successive null-hypothesis tests. (See Supplementary Material Sec. 5.) For each mutant *i* described by the overabundance model, the TFNseq experiment measures a growth arrest time *T*_*i*_ and the corresponding target protein overabundance:

**FIG. 3.**
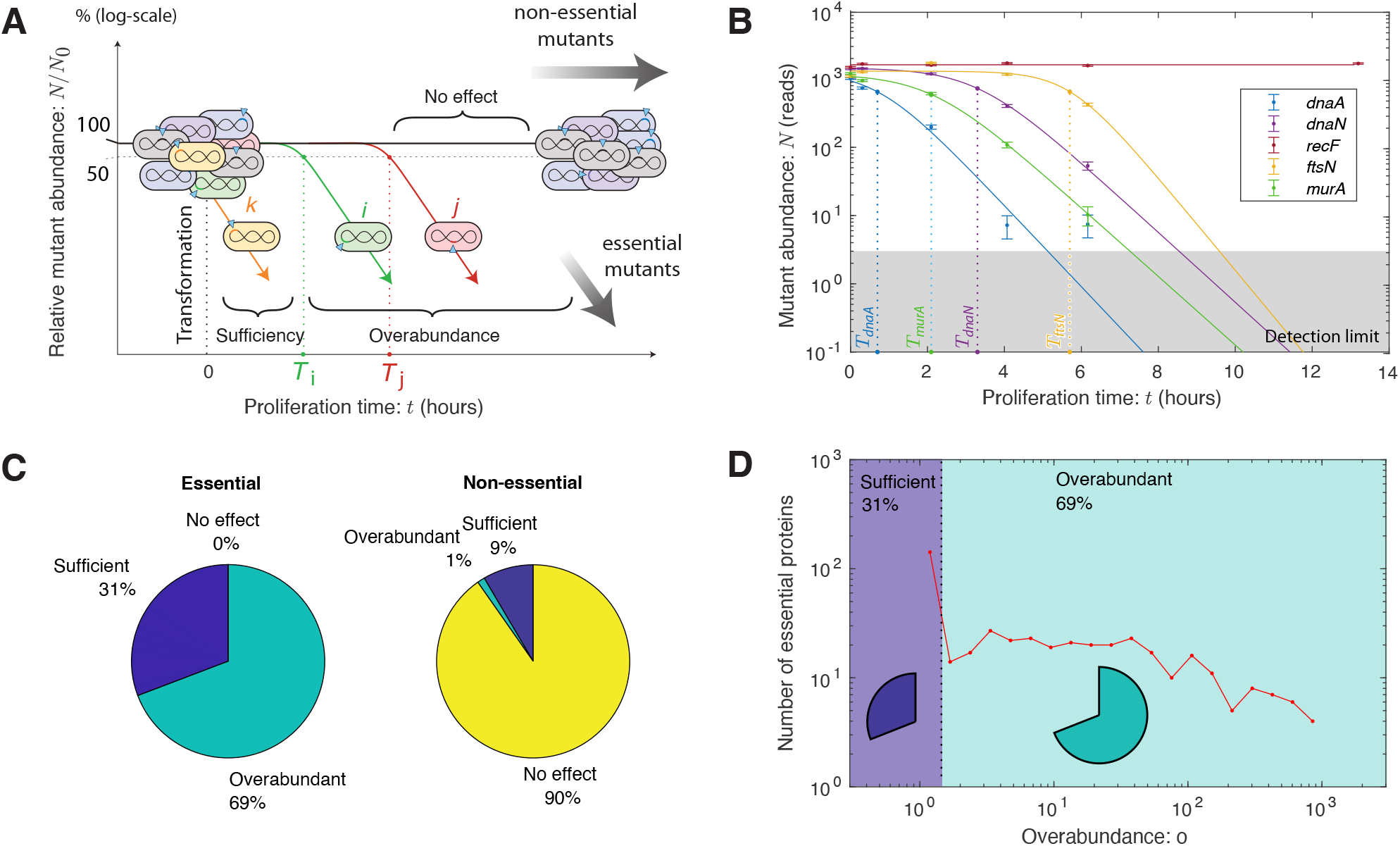
A proteome-wide analysis of protein overabundance. Panel A: TFNseq schematic. A poly-clonal library of knockout mutants is generated by the transformation of ADP1 with DNA mutagenized by transposon insertions. The library is proliferated on selective media and sequential fractions are collected. The relative-abundance trajectories of mutants are determined by mapping transposon insertion sites by sequencing. The arrest time parameter *T*_*i*_ corresponds to the time at which the growth rate changes from wild-type to slow growth for mutant *i*. The arrest times are represented as dotted lines, corresponding to the time at which the relative abundance falls to 50% its initial value. (See Supplementary Material Sec. 4 C.) **Panel B: TFNseq-trajectory analyses for five mutant strains**. Each mutant trajectory is well fit by one of the three trajectory models. As expected, the no-effect model is selected for the non-essential gene *recF*. For the other four essential genes, the overabundance model is selected. The dotted line represents the arrest time for each mutant. **Panel C: Proteome-wide analysis of proliferation dynamics**. For genes classified as essential, 31% were best fit by the sufficiency-dynamics model while 69% were best fit by the overabundance trajectory model. For genes classified as non-essential, 90% were best fit by the no-effect model, while 10% showed a detectable reduction in growth rate. **Panel D: Overabundance varies by orders of magnitude between essential proteins**. The protein overabundance is inferred from the arrest time using Eq. 4. Sufficient expression genes have overabundance *o* = 1, while over-abundant genes vary from *o >* 1 to very large overabundance (*o >* 100).

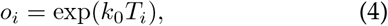

where *k*_0_ is the wild-type growth rate. Two replicate experiments lead to comparable overabundance estimates. (See Supplementary Material Sec. 5.)

To test the consistency of this TFNseq approach with imaging-based knockout-depletion measurements, we focused first on the analysis of the mutants *dnaA, dnaN, ftsN*, and *murA*. As shown in Fig. 3B, the trajectories for *dnaA, murA, ftsN*, and *dnaN* show an unambiguous steplike change in growth dynamics: The no-effect trajectory model (null hypothesis) are rejected with p-values that are below machine precision, and the sufficiency trajectory model is also rejected with *p <* 10^−4^ for all genes. In Tab. I, we compare protein overabundances determined by imaging- and sequencing-based approaches. These numbers are qualitatively consistent. For instance, the single-cell analysis of *dnaA* mutant shows a nearly immediate phenotype by imaging (*i*.*e*. cell filamentation). (See Supplementary Sec. 1 E.) Likewise, the TFNseq-approach finds an overabundance of 1.0, meaning that protein expression is sufficiency. On the other hand, all three of the other mutants (*murA, ftsN*, and *dnaN*) are found to have very large overabundances, and are roughly comparable. Finally, a representative non-essential gene (*e*.*g. recF*) shows no effect. These results support the use of the TFNseq approach to analyze protein overabundance genome wide.

### Many essential proteins have vast overabundance

To determine the protein overabundance genome-wide, we analyzed the knockout-depletion trajectories for all genes in *A. baylyi*. (See Fig. 3BCD.) Our analysis showed that the vast majority (90%) of genes annotated as non-essential were classified as having *no effect* and 10% of non-essential genes had measurable growth defects. (Replicate experiments were used to estimate the error in the protein overabundance. See Supplementary Material Fig. S13.) The most severe growth defect in non-essential annotated genes were observed for the genes *gshA* and *rplI*.

For the essential gene analysis, we defined essential genes conservatively, demanding that they be classified essential by both TFNseq [18] and single-gene deletion experiments (432 genes) [27]. All essential mutants were observed to have growth defects, as anticipated; however, only 31% of essential proteins were classified as *sufficient*, corresponding to an immediate change in growth rate. Notable genes in this category include ribosomal proteins RpsQ and RpsE, ribonucleotide reductase subunits NrdA and NrdB, and ATP synthase subunits AtpA and AtpD. However, as predicted by the RLTO model, the majority of essential proteins (69%), were classified as *overabundant*, meaning that they required significant dilution before a growth rate change was detected. Fig. 3D shows a histogram of essential gene overabundances.

### Low-transcription genes are highly overabundant

To understand the overall significance of overabundance in a typical biological process, we determined the median essential protein overabundance: 7-fold. To understand the significance of overabundance from the perspective of the metabolic load, we also determine the mean protein overabundance, weighted by the expression level: 1.6-fold. These two superficially-conflicting statistics emphasize a key predicted regulatory principle: Over-abundance is high for low-abundance proteins; however, it is close to unity for the high-abundance proteins, which constitute the dominant contribution to the metabolic load.

To explicitly test the predicted relation between protein expression and overabundance, we quantified transcription by measuring the relative abundance of mRNA messages by RNA-Seq for exponentially growing *A. baylyi* cells. (See Supplementary Material Sec. 6.) We estimated transciption, in mRNA transcripts per gene per cell cycle (*message number*) for each essential gene. (See Supplementary Material Sec. 6 B.) Fig. 4A compares transcription and overabundance for all essential genes with the prediction of the RLTO model. To quantitatively capture the trend, we applied two different binning approaches to compare the data cloud to the RLTO model predictions. As predicted, the data show a clear trend of decreasing overabundance with increasing message number. With very few exceptions, high-expression genes have extremely low overabundance. At the other extreme, low expression genes typically have large to very large overabundance as shown by the sharp up-turn of the purple curve as the transcription level approaches the one-message-rule threshold, a lower threshold on essnetial-gene transcription that we recently proposed [11].

**FIG. 4.**
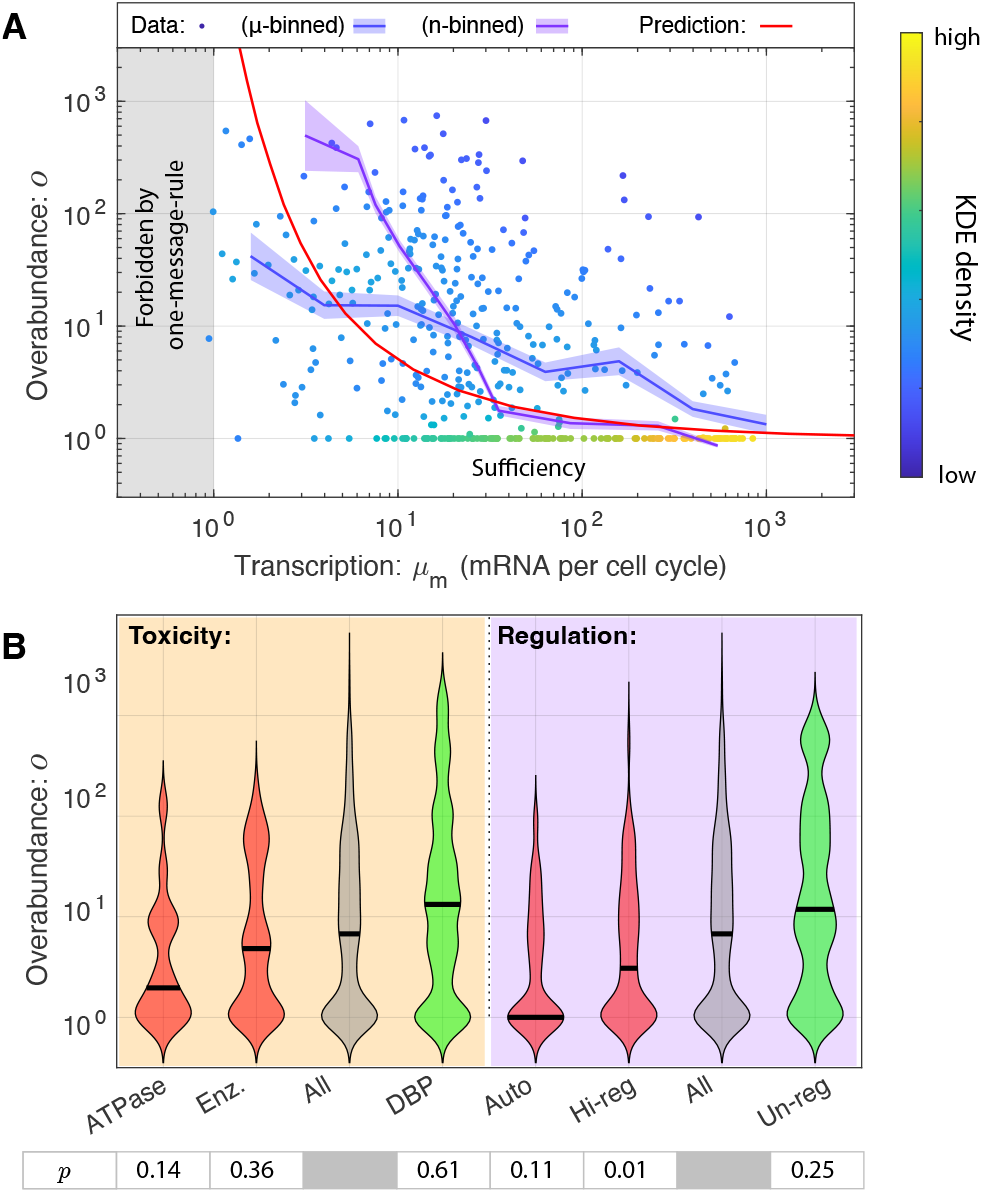
Panel A: Overabundance is large for lowexpression essential proteins. The measured transcription-overabundance pairs are shown for all essential genes. (The density of data points was estimated with a KDE.) To analyze the trend, this data was binned using two complementary approaches (blue and purple) (See Supplementary Material Sec. 5 C.) The RLTO model (red) predicts that overabundance grows rapidly as the transcription level is reduced. The RLTO model (red) qualitatively captures the trend of the data; however, it appears to underestimate the measured overabundance for intermediate expression genes. The RLTO model predicts that essential genes cannot be expressed below the level of one mRNA per cell cycle (one-message-rule) [11]. **Panel B: Toxicity and regulation are determinants of over-abundance**. We compared the overabundance measurements for six essential gene subgroups to determine whether toxicity and regulation could affect overabundance. Red groups were predicted to decrease overabundance while green groups were expected to increase it. The p-values for the consistency of each distribution with the all gene group is shown below each category and is generated using the Kolmogorov-Smirnov test [28]. As hypothesized, the data is consistent with both toxicity and regulation decreasing overabundance.

### Limitations of knockout-depletion experiments

In spite of the success of the RLTO model in predicting the genomic-scale overabundance trend, there are many significant outliers from this prediction. In considering their significance, it is important to emphasize the flaws both with the knockout-depletion experiments, as well as the RLTO model. With respect to the experiments, the mechanism of growth arrest plays an important role in determining which growth metric most accurately determines the arrest time. Consider the three arrest times measured for the septation-related essential gene *ftsN* in Tab. I. Due to the absence of strict cell-cycle checkpoints in the bacterial cell, the arrest of the septation process does not immediately arrest cell elongation and replication [29]. Growth arrest is therefore detected first by the cell-number metric, directly dependent on septation, and later in the other two metrics. The genomewide TFNseq-approach measures replication-based arrest and is therefore expected to be most precise for replication genes and overestimate the arrest time for other processes. In spite of this limitation, the TFNseq approach is tractable at a genomic-scale and is accurate enough to capture transcription-based trend predicted by the RLTO model (Fig. 4A).

### Other determinants of overabundance

In addition to these experimental limitations, the RLTO model itself relies on a number of important simplifications. Several authors have already focused on the significance of protein function which appears to play a significant role in shaping the fitness landscape [9, 18, 30]. Here, we will focus on exploring two specific function-related hypotheses that affect RLTO model predictions [11]: the role of gene regulation and protein toxicity in determining overabundance.

### Tightly regulated genes have low overabundance

A key assumption in the RLTO model is that gene expression noise is a consequence of the number of mRNA only and is otherwise independent of regulation [11]. Precise control of expression could lead to a reduction in the optimal overabundance. To explore the regulatory hypothesis, we generated three lists of essential genes: auto-regulatory, highly-regulated (top 10% of genes ranked by number of regulators), and unregulated. If regulation can obviate the need for over-abundance, we would expect lower median overabundances in both regulated groups and potentially higher overabundances for the un-regulated group relative to all essential genes. Consistent with this hypothesis, we find that the median overabundance for auto-regulatory genes is 1-fold and for highly-regulated, 3-fold, compared with 7-fold for all essential genes, and 12-fold for un-regulated genes, supporting the hypothesis that tight regulation could reduce the need for overabundance. (See Fig. 4B.)

### Proteins with functional potential for toxicity have low overabundance

A second key assumption in the RLTO model is that the metabolic cost of transcription and translation are the dominant fitness costs of protein overabundance (*i*.*e*. there is no toxicity) [11]. To explore the potential role of toxicity, we generated groups of essential ATPases and enzymes, hypothesizing that these proteins would have higher cost due to excessive activity when overabundant, and a group of DNA-Binding Proteins (DBP), which we hypothesized would have low cost when overabundant. We find that the median overabundance for ATPase genes is 2-fold, and for enzymes more generally 5-fold, compared to 7-fold for all essential genes and 13-fold for DBP. These results are consistent with the hypothesis that toxicity, and in particular ATPase activity, is also a key determinant of overabundance. (See Fig. 4B.)

## DISCUSSION

### The shape of the fitness landscape

Despite some largescale measurements [8, 9, 30–32], fundamental questions remain about the structure of the fitness landscape and its rationale [7]. Our genome-wide measurements reveal that most (69%) essential proteins are consistent with a step-like transition between wild-type and arrested growth below a critical threshold protein abundance. These step-like transition are qualitatively consistent with asymmetric landscapes that have been observed previously which transition more gradually from arrest of slow growth to a relatively flat plateau (*e*.*g*. [3, 30, 31]). Does the detailed form of the fitness landscape affect RLTO predictions? It is important to emphasize that optimality of overabundance does not depend on the detailed mathematical form of the fitness landscape: It is the strong asymmetry of that landscape that is required to predict protein overabundance [11].

How do we reconcile the sharpness of the transition between growth and no-growth in the single-cell image-based analysis with previous measurements that show a more complex transition? An intriguing possibility is that cell-to-cell variation in protein abundance may significantly obscure the underlying landscape. In imaging-based experiments, once protein expression ceases, the remaining proteins are partitioned between daughter cells with very low noise [22]. A sharp transition is therefore expected in the progeny that begin from a common pool of protein in a single progenitor cell (Fig. 1A). We hypothesize that the observed progenitor-to-progenitor variation in the arrest time is due to differences in the single-cell protein abundances in the progenitor cells (Fig. 2C). As previously been discussed [16], steady-state CRISPRi-based protein depletion may increase cell-to-cell variation in protein expression beyond the existing endogenous expression noise, further smoothing the underlying fitness landscape. To our knowledge, there has yet to be any consistent analysis of noise in CRISPRi silencing in bacterial cells, even though this phenomenon could clearly limit the resolution of steady-state CRISPRi-based experiments. That said, the knockout-depletion approach or analogous CRISPRi-based dilution approaches [9], is not a steady-state measurement of cell growth and therefore this dilution-based approach to measuring the fitness landscape may introduce its own systematic limitations.

### The rationale for a threshold abundance

The observed threshold-like dependence can be rationalized in terms of chemical kinetics: Changing the abundance of a reactant in a multi-component reaction will only change the rate if the reactant is rate-limiting [11, 33]. As the abundance of the rate limiting reactant *i* is increased, the reaction rate also increases until another reactant becomes rate-limiting, after which the rate saturates with respect to *i*, giving rise to a threshold. (See Fig. 5.) We therefore hypothesize that the threshold-like behavior observed in the fitness landscape is the consequence of nearly all proteins being expressed in excess of the rate-limiting concentration. Consistent with this picture, we explicitly demonstrate protein function (*i*.*e*. replication) is robust to an order-of-magnitude depletion of replisome protein DnaN; however, for most proteins, we must infer this picture from the growth rate.

**FIG. 5.**
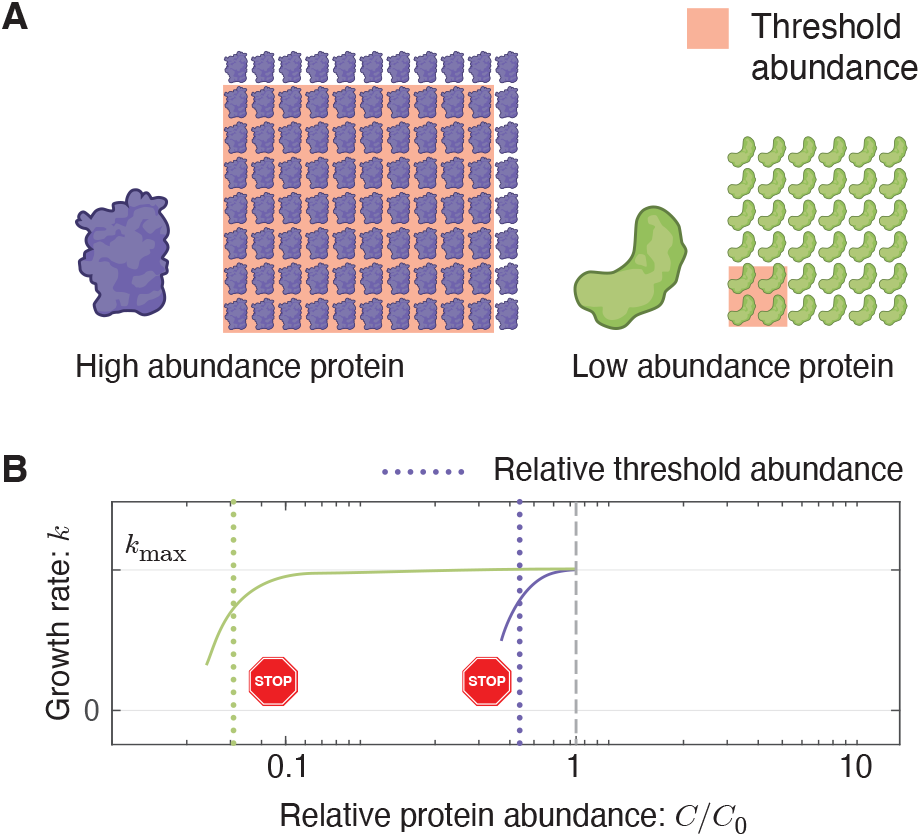
How rate-limited kinetics shapes the fitness landscape. Panel A: Protein abundance and threshold. Two essential protein species with different abundances are pictured schematically. The threshold abundance at which each protein becomes limiting is represented by the pink square and the total cellular abundance is represented by the protein array. **Panel B: Emergent fitness landscape**. A schematic model of the growth rate versus relative protein abundance is shown for the two protein species. The RLTO model predicts that low-abundance proteins (green) have high overabundance, which leads to significant insensitivity to protein depletion. High-abundance protein (purple) are predicted to have small overabundance leads to high sensitivity to protein dilution. The growth rate rapidly decreases with concentration once a species becomes limiting.

### The rationale for overabundance

Rate-limiting kinetics does not in itself predict vast protein overabundance. The RLTO model predicts that this feature of the fitness landscape is a consequence of a balance between (i) the metabolic cost of protein expression, which favors minimizing protein abundance, and (ii) robustness to the noise in gene expression [34, 35]. The model predicts expression-dependent protein overabundance: large overabundance for low-abundance proteins and small overabundance for high-abundance proteins due to the highly asymmetric fitness trade-off: the metabolic cost of overabundance for low-expression genes is small compared with the failure of essential processes [11]. We show that this signature prediction is observed (Fig. 4A).

### Environmental fluctuations

The overabundance predicted by the RLTO model hedges against noise, *i*.*e*. the cell-to-cell variation in gene expression; however, it has also been proposed that overabundance might result from hedging against a second form of fluctuations, environmental fluctuations, in which the cell transition from one carbon sources to another or from good to poor growth conditions [36, 37]. We do not reject this hypothesis outright, especially in the context of genes involved in metabolism; however, the mechanism we propose is much more generic and predicts overabundance should be optimal even for low-expression proteins with little direct connection to metabolism (*e*.*g*. replication, cell-wall synthesis, *etc*.).

### Is protein overabundance conserved?

To what extent is the overabundance of essential genes a conserved mechanism from bacteria, to single-cell eukaryotes, to multicellular organisms? If overabundance were specific to *A. baylyi*, we would expect to dramatic differences between the *A. baylyi* and *E. coli* transcriptomes for low-expression essential-gene homologues. This is not observed (Supplementary Material Fig. S11.) As we emphasized above, CRISPRi protein depletions in a wide range of model organisms appear to be qualitatively consistent with the overabundance hypothesis in that large-factor depletions are often required to generate a phenotype [8, 12–15]. Furthermore, we have demonstrated elsewhere that the RLTO model also predicts two other principles of central dogma function (the onemessage-rule and load balancing in protein expression) that are observed in eukaryotic cells [11]. We therefore expect overabundance strategy to be conserved in all organisms for low-expression genes.

### Are non-essential proteins overabundant?

We have focused our analysis on essential genes in the model organism *A. baylyi* and demonstrated that most essential proteins are overabundant. To what extent is this mechanism generic to non-essential proteins? Several arguments support a generic applicability to non-essential genes. Our modeling suggests asymmetry rather than explicit growth arrest is the mathematical rationale for the optimality of overabundance [11]. We therefore predict that all proteins that increase cell fitness, not just essential proteins, will be overabundant. In addition, it is important to emphasize that the annotation of genes as *essential* is contextual. For instance, for *E. coli* proliferation on lactose, the gene *lacZ* is essential, although non-essential for other carbon sources. As a result, we predict that when induced, LacZ should be overabundant, consistent with observation [36]. Finally, the RLTO model also correctly predicts the balance between transcription and translation for all genes, not just essential genes, in eukaryotic cells, suggesting that it should generalize to nonessential genes as well [11].

### Biological implications

Many important proposals have been made about the biological implications of noise [38]. Our work reveals that noise acts to inflate the optimal expression levels of low-expression proteins and, as a result, significantly increases the metabolic budget for protein, which constitutes 50-60% of the dry mass of the cell [4]. We believe this increased protein budget has cellular-scale implications. For instance, in stress response and stationary phase, the presence of a significant reservoir of overabundant protein provides critical resources, via protein catabolism, to facilitate the adaptation to changing conditions [39, 40]. Protein overabundance may have important implications for individual biological processes as well, including determining which proteins and cellular processes make attractive targets for small molecule inhibitors (*e*.*g*. antibiotics) [32]. Since overabundance defines the fold-depletion in protein activity required to achieve growth arrest, high-overabundance proteins are predicted to be extremely difficult targets for inhibition.

## Conclusion

By combining imaging-, genomic-, and modeling-based approaches, we provide a both a quantitative measurement of the fitness landscape for all essential proteins as well as a clear qualitative and conceptual understanding of the rationale for the observed fitness landscape. The RLTO model fundamentally reshapes our understanding of the rationale for protein abundance. The model predicts, and experiments confirm, that low-abundance proteins are expressed in vast excess of what is required for growth. Despite the limitations of the experiments, the predicted trend is clearly resolved both at a genomic-scale, using sequencing-based approaches, as well as at the single-cell scale, as observed by microscopy. The rationale for the overabundance strategy is intuitive: Growth requires the robust expression of hundreds of distinct proteins. The cell contends with this extraordinary complex regulatory challenge by keeping all but the highest-abundance proteins in vast excess.

## Data availability

We include source data files and sequencing data from RNA-Seq experiments to quantify transcription levels. NCBI Sequence Read Archive, https://www.ncbi.nlm.nih.gov/sra (accession no. PRJNA1173237).

## Acknowledgments

The authors would like to thank B. Traxler, A. Nourmohammad, J. Mougous, M. Cosentino-Lagomarsino,. van Teeffelen, C. Manoil,L. Gallagher, J. Bailey, J. Mannik, G. Chure and S. Murray. H.J.C., T.W.L., D.H., and P.A.W. were supported by NIH grant R01-GM128191 and NSF grant GR046955.K.J.C. was supported by the Molecular Biophysics Training Program (NIH grant T32GM008268). W.R.W. was supported by NIH grant R01-AI150041.

## Author contributions

H.J.C., T.W.L., K.J.C., D.H., and P.A.W. conceived the research. H.J.C., W.R.W., and P.A.W. performed the experiments. H.J.C., T.W.L., K.J.C., and P.A.W. performed the analysis. H.J.C., T.W.L., D.H., and P.A.W. wrote the paper.

## Competing interests

The authors declare no competing interests.

### Supplementary Materials

Supplementary text, Materials and Methods

Figs. S1 to S11.

Tables S1 to S3.

References 35 to 56.

Data files S1 to S9.

Movies S1 to S12.

## Supplementary material

## 1. RESULTS: IMAGING-BASED KNOCKOUT-DEPLETION EXPERIMENTS

### A. Some progenitors have heterogenic progeny

Heterogenic progenitors are progenitor cells that are observed to have progeny with two distinct heritable phenotypes: the Km^R^ knockout phenotypes and the Km^S^ wild-type phenotype. For instance, in the Δ*murA* knockout-depletion experiments, progenitors were observed with one daughter whose progeny proliferated for multiple generations on Km^+^ media before lysing, the knockout phenotype, and whose other daughters proliferated for a short period but maintained wild-type morphology. The maintenance of the wild-type morphology suggested that the cells were *murA*^+^ Km^S^. How were these cells able to proliferate while other Km^S^ cells immediately arrested?

We hypothesize that since both cells had the same progenitor, recombination occurred in the mother cell, after the *murA* gene was replicated, leading to one wildtype chromosome and one Δ*murA* chromosome. The transient growth of the wild-type cells was the result of overabundance of the *kan* gene product APH(3’)II being expressed before cell division in the original mother cell.

Heterogenic progenitor cells appeared frequently for *dnaN* knockout-depletion experiments, presumably because of the location of *dnaN* in the immediate vicinity of the origin, resulting in early replication. In these experiments, an additional test of the heterogenic progenitor hypothesis was possible due to the fluorescent labeling of the target protein. Cells that arrested early with the wild-type morphology showed no protein depletion; whereas cells that displayed the mutant phenotype (filamentation) showed depleted YPet-DnaN levels.

**FIG S1.**
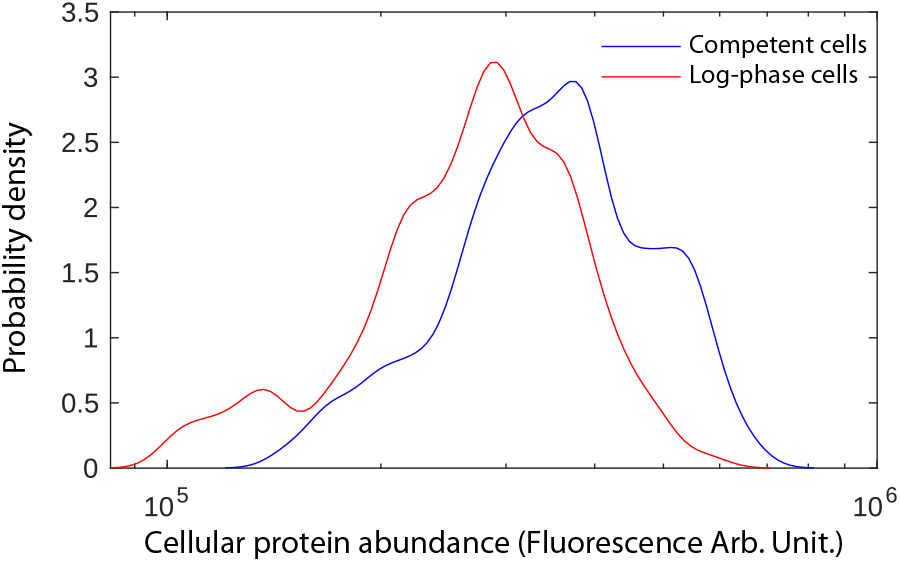
Panel A: Protein abundance in competent cells are comparable to log phase. We characterized YPet-DnaN abundance in transformed cells *before protein dilution* (at the end of out growth after transformation, *N* = 115 cells) and wild-type cells (*N* = 157 cells). The distribution of cellular protein abundances are shown above, with the median abundance of YPet-DnaN 27% higher in transformed cells.

### B. Transformed cells start with log-phase protein abundances

The experimental design of knockout-depletion assays (Fig. 1A) fails if cells do not start initially with wildtype protein abundances. The question is necessitated by the transformation protocol: Cells begin in stationary phase and are mixed with DNA knockout cassette. The cells are then propograted on non-selective media for 2.5-3 h. (See Sec. 4 A 1.)

To test the design, we characterized YPet-DnaN abundance in transformed cells (at the end of out growth after transformation) and wild-type cells. We found that the protien expression levels in the two states were comparable, with the median abundance of YPet-DnaN 27% higher in transformed cells. (See Fig. S1.) This comaparable expression level is to be expected since the competent cells are grown for more than two generations the same media as log-phase cells.

### C. Analyses of dilution rate of YPet-DnaN

The experimental design of knockout-depletion assays (Fig. 1A) fails if there is a delay before either the transformation or recombination processes. If the DNA target sequence is not rapidly degraded and expression continues, transient growth could be the consequence of this continuing expression. To test for this possibility, we determined the dilution rate of the fluorescently-labeled beta clamp (YPet-DnaN) in the YdnaN strain after knockout. If protein is depleted by dilution, the fluorescence intensity should decay with the same rate as the cell progeny area grows (a proxy for volume) grows (Eq. 1). In our knockout experiment, we identified four transformed progenitor cells that were isolated enough to analyze the fluorescence intensity during cell proliferation. We determined both the dilution and growth rate and compared them. (See Fig. S2A.) In each case, the rates were consistent. (See Fig. S2B.)

**FIG S2.**
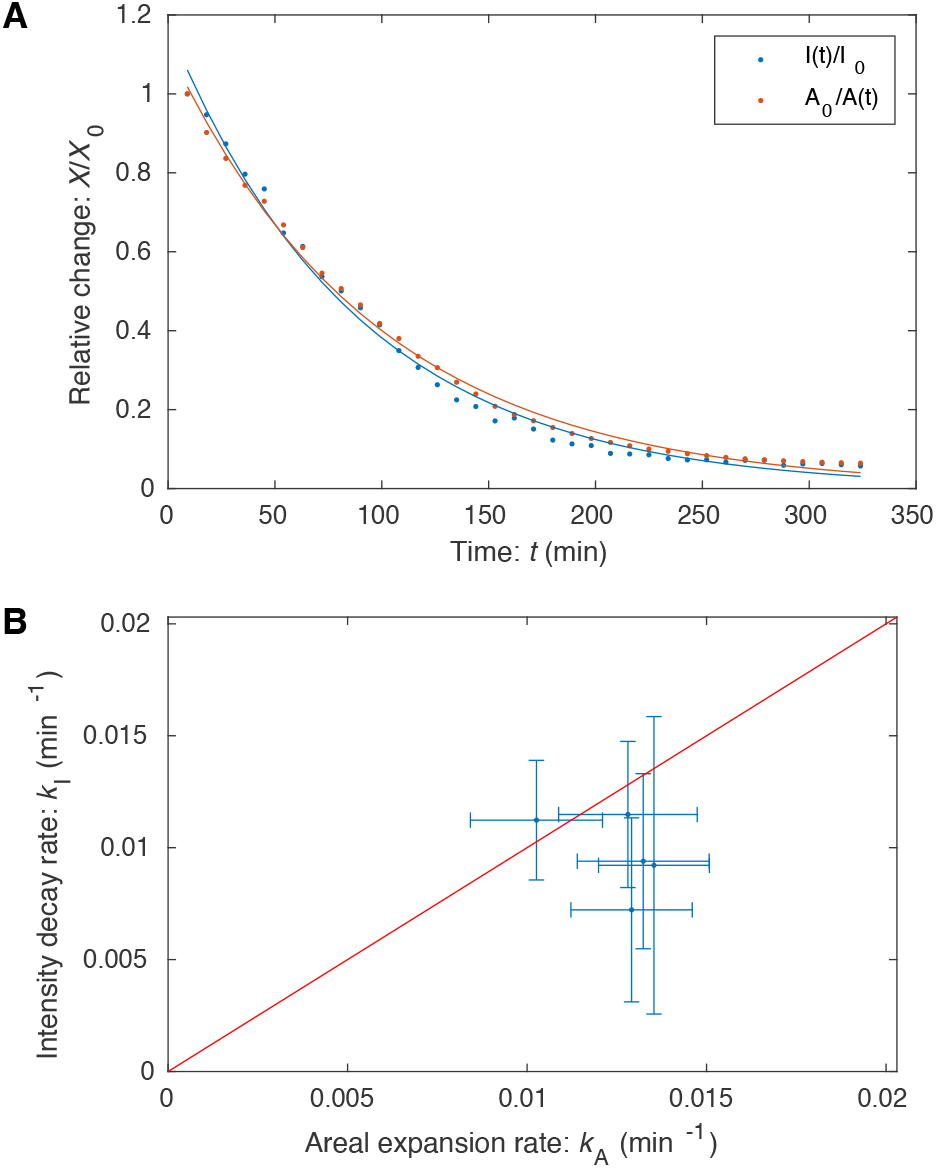
Panel A: Protein abundance and growth in a YPet-DnaN cell after knockout. The background-subtracted average intensity and inverse area were fit to exponential functions to determine their respective rates of decay for each micro colony. **Panel B: Dilution versus growth rate**. The fluorescence decay rate *kI* is compared to the areal growth rate *k*_*A*_ for four microcolonies. The experimental design assumes that these rate are equal. The observed rates are consistent with this assumption. The sensitivity of this experiment was limited by the fluorescence of neighboring wild-type cells.

This experiment suffers from important limitations. Transformants nearly always wild-type cells, as described in the last section, and fluorescence of these neighboring cells limits the sensitivity of the assay at late times due to the diffuse background generated by the brightly fluorescent wild-type cells.

### D. Wild-type imaging-based analyses

We analyzed two different strains with wild-type growth phenotypes: wild-type cells (*Acinetobacter baylyi* ADP1) and ACIA0320-0321∷*kan*.

*IS∷kan*. To generate a reference wild-type growth phenotype, we choose a non-essential gene with no reported phenotype, genes ACIA0320-0321, corresponding to an IS element. The deletion was performed on the YdnaN strain, which shows no growth phenotype under the experimental conditions. We will abbreviate this strain Δ*IS*. We constructed this deletion and measured its growth relative to wild-type on Km^-^ media, and no growth phenotype was observed. However, even though this strain can be stably maintained (since ACIA0320-0321 is non-essential), we transformed this cassette using the same protocol in knockout-depletion experiments. As expected, a comparable number of transformants were observed using this construct to those targeting essential genes.

**TABLE S1.**
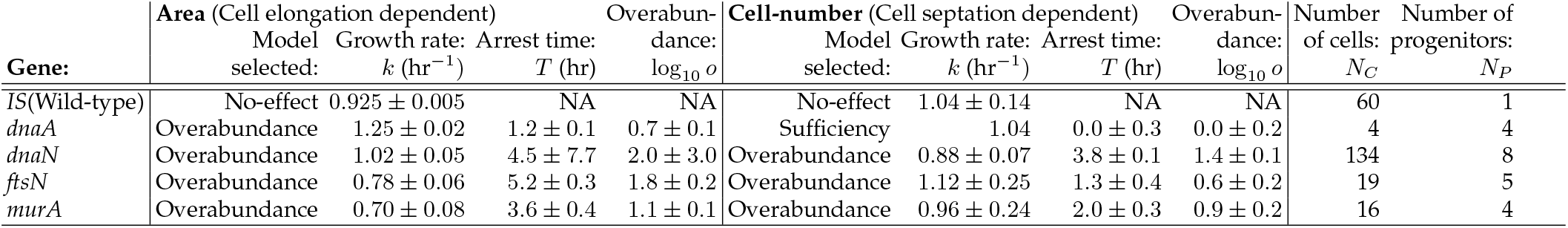
Detailed results from fitting imaging-based knockout-depletion experiments. The table summarizes the analysis of cell proliferation by two complementary metrics: area and cell-number analyses. These two metrics depend on distinct cellular processes: Growth in cell area is dependent on cell elongation, whereas the proliferation of cell number is dependent on the septation process. We give two metrics for sample size: the number of progenitors (*N*_*P*_) and the total number of cells analyzed (*N*_*C*_), corresponding to progenitor and progeny. The estimated standard error is provided for parameter fits.

A typical transformant from a knockout-depletion experiment targeting *IS* is shown in Fig. S3 for which six generations of growth are captured. Both the areal (cell-elongation-dependent) and cell-number (septation-dependent) analyses are consistent with the null hypothesis, the *No-effect model*, as expected. The growth rate was observed to be *k* = 0.925±0.005 hr^−1^ for the areal analysis and *k* = 1.04 ±0.14 hr^−1^ for the cell-number analysis.

#### Qualitative phenomenology

A typical knockout-depletion experiment is shown in Fig. S3. Panel A shows a frame mosaic. The cells in this dataset show the log-phase growth phenotype of wild-type cells. Both cell number and area show exponential growth. The step-like growth of the cell number reflects the desynchronization of cell division events of the ancestors for a single progenitor.

#### Quantitative analysis

The null hypothesis (*Sufficiency model*) was rejected in favor of the No-effect model for both the area and cell-number analysis (both p-values under machine precision). The growth rate was observed to be *k* = 1.04 ± 0.14 hr^−1^ for the areal analysis and *k* = 0.925 ± 0.005 hr^−1^ for the cell-number analysis.

### E. *dnaA* imaging-based analysis

#### Annotated gene function

DnaA is an essential regulator of the cell cycle and DNA replication initiation in particular.

#### Qualitative phenomenology

A typical knockout-depletion experiment is shown in Fig. S4. Panel A shows a frame mosaic. The cells in this dataset show the onset of the phenotype, cell filamentation, without undergoing significant growth-induced protein dilution. As a result, the cell number, shown in Panel B, is constant since no divisions are observed. However, as shown in Panel C, cell elongation continues for roughly 100 min before it begins to arrest. We interpret the metric that shows the earliest arrest to define the overabundance. In this case, since septation is not observed again after transformation, DnaA abundance is consistent with the Sufficiency model.

#### Quantitative analysis

The null hypothesis (*Sufficiency model*) was rejected in favor of the *Overabundance model* (p-value under machine precision) for the areal analysis. The initial growth rate was observed to be *k* =1.25 ±0.02 hr^−1^ with an arrest time of *T* = 1.24± 0.10 hr.In case of the cell number analysis, we fail to reject the null hypothesis (*Sufficiency model*), indicating that there is no statistical significance to support the alternative hypothesis *No-effect model* (p = 1.0). We used the Δ*IS* wild-type growth rate (*k* = 0.925 ± 0.005 hr^−1^) to fit the arrest time: *T* = 0.0 ± 0.3 hr.

### F. *dnaN* imaging-based analysis

#### Annotated gene function

The gene product of *dnaN* is the *β* sliding clamp (DnaN), which is an essential component of the replisome complex.

#### Qualitative phenomenology

A typical knockout-depletion experiment is shown in Fig. S5. Panel A shows a frame mosaic. The cells in this dataset show the onset of the phenotype, cell filamentation, at about 220 min, after multiple rounds of cell division. As a result, the cell number, shown in Panel B, plateaus shortly after the filamentation is observed since the filamentation is a consequence of the failure of the cells to efficiently septate. However, as shown in Panel C, cell elongation continues, although slowing slightly, throughout the experiment. In this case, since arrest is observed first with respect to septation, we use the arrest of this process to define overabundance.

**FIG S3.**
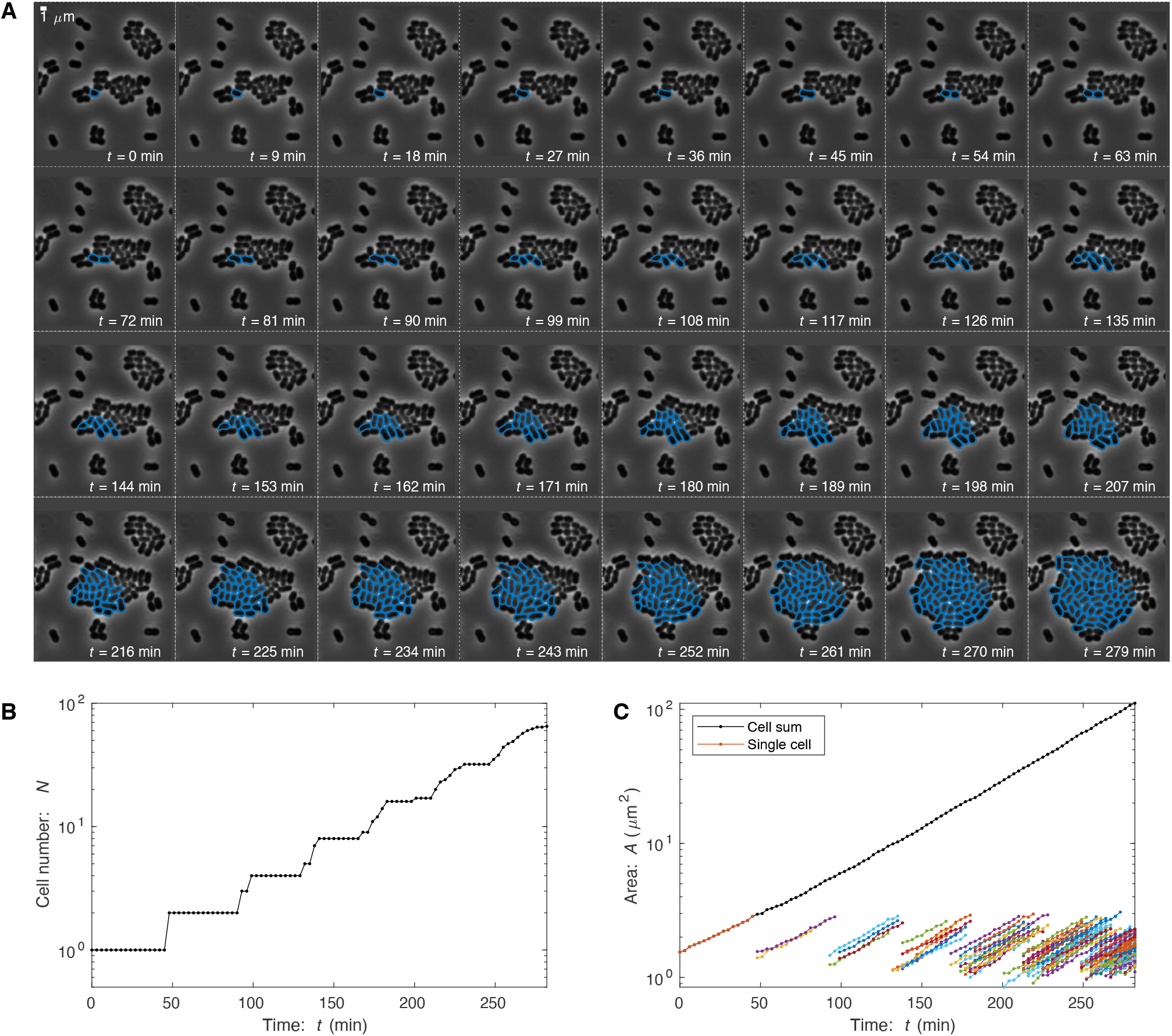
Knockout-depletion experiment: IS element (Non-essential). Panel A: Frame mosaic. In the knockout-depletion experiment, the majority of cells are not transformed and immediately arrest on media supplemented with kanamycin. The lone transformant (*IS*∷*kan* (Km^*R*^), blue) proliferates normally. Cells were segmented using OmniSegger for quantitative analysis. **Panel B: Cell number**. The number of transformant progeny as a function of time. **Panel C: Progeny area**. Total progeny-cell area as a function of time. Total cell area is plotted with the black-dotted line, while individual cell areas are plotted with color.

#### Quantitative analysis

The null hypothesis (*Sufficiency model*) was rejected for both the area (*p* = 8.9 ×10^−140^) and cell-number analysis (*p* = 6.0× 10^−19^). The initial growth rate was observed to be *k* = 1.02± 0.05 hr^−1^ with an arrest time of *T* = 4.5 ±7.7 hr for the areal analysis. For cell-number analysis, the initial growth rate was observed to be *k* = 0.88 ± 0.07 hr^−1^ with an arrest time of *T* = 3.8 ± 0.1 hr.

### G. *ftsN* imaging-based analysis

#### Annotated gene function

The gene product of *ftsN* is essential cell division protein FtsN.

#### Qualitative phenomenology

A typical knockout-depletion experiment is shown in Fig. S6. Panel A shows a frame mosaic. The cells in this dataset show the onset of the phenotype: the failure to septate, at roughly 150 minutes, after several rounds of division. As a result, the cell number, shown in Panel B, plateaus shortly after 150 min as a consequence of the failure of the cells to efficiently septate. However, as shown in Panel C, cell elongation continues, although slowing slightly, to roughly 220 min.

**FIG S4.**
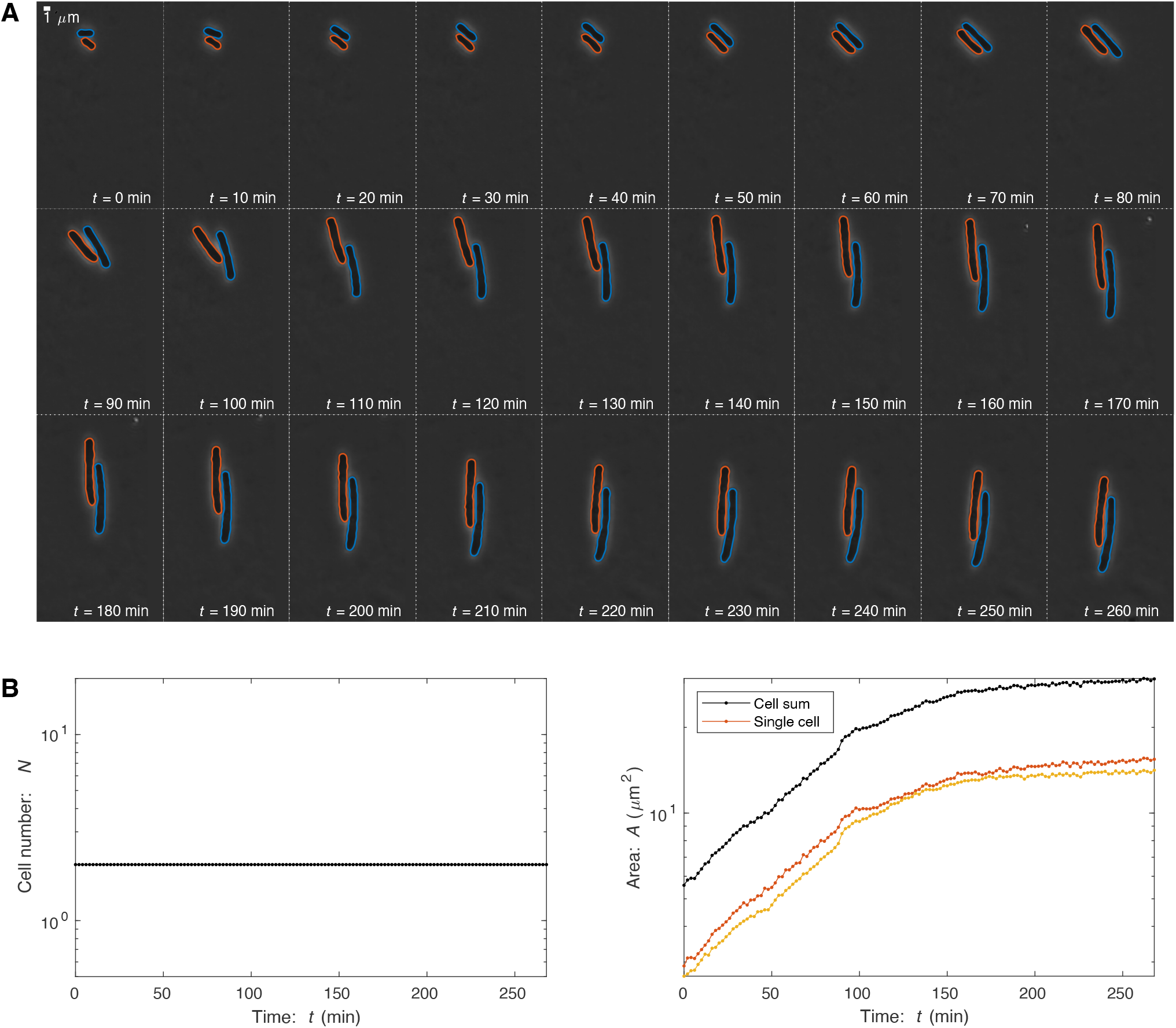
Knockout-depletion experiment: Δ*dnaA*. Panel A: Frame mosaic. Two transformants (*dnaA*∷*kan*(Km^*R*^), blue, orange) proliferate. DnaA is an essential regulator of replication initiation. Its depletion leads to a failure of the chromosome to replicate, and therefore results in cell filamentation. Cells were segmented using OmniSegger for quantitative analysis. **Panel B: Cell number**. The number of transformant progeny as a function of time. After transformation, cells fail to divide, consistent with DnaA expression being sufficient rather than overabundant. **Panel C: Progeny area**. Total progeny-cell area as a function of time. In spite of the arrest of septation/division, cell areal elongation persists for roughly 120 minutes.

#### Quantitative analysis

The null hypothesis (*Sufficiency model*) was rejected for both the area (p-value under machine precision) and cell-number analysis (*p* = 1.7 ×10^−7^). The initial growth rate was observed to be *k* =0.78 ±0.06 hr^−1^ with an arrest time of *T* = 5.2 ±0.3 hr for the areal analysis. For cell-number analysis, the initial growth rate was observed to be *k* = 1.12 ± 0.25 hr^−1^with an arrest time of *T* = 1.3 ± 0.4 hr.

### H. *murA* imaging-based analysis

#### Annotated gene function

The gene product of *murA* is UDP-N-acetylglucosamine 1-carboxyvinyltransferase, an essential protein in synthesizing the precursors of cell wall synthesis.

#### Qualitative phenomenology

A typical knockout-depletion experiment is shown in Fig. S7. Panel A shows a frame mosaic. The cells in this dataset show the onset of the phenotype: the loss of cell wall integrity, and therefore first the loss of wild-type cell morphology and then cell lysis. Cells begin to lose their wild-type morphology at roughly 120 min, after multiple rounds of cell division. As a result, the cell number, shown in Panel B,plateaus shortly after 150 min as a consequence of the failure of the cells to efficiently septate. However, as shown in Panel C, cell elongation continues, although slowing slightly, to roughly 200 min.

**FIG S5.**
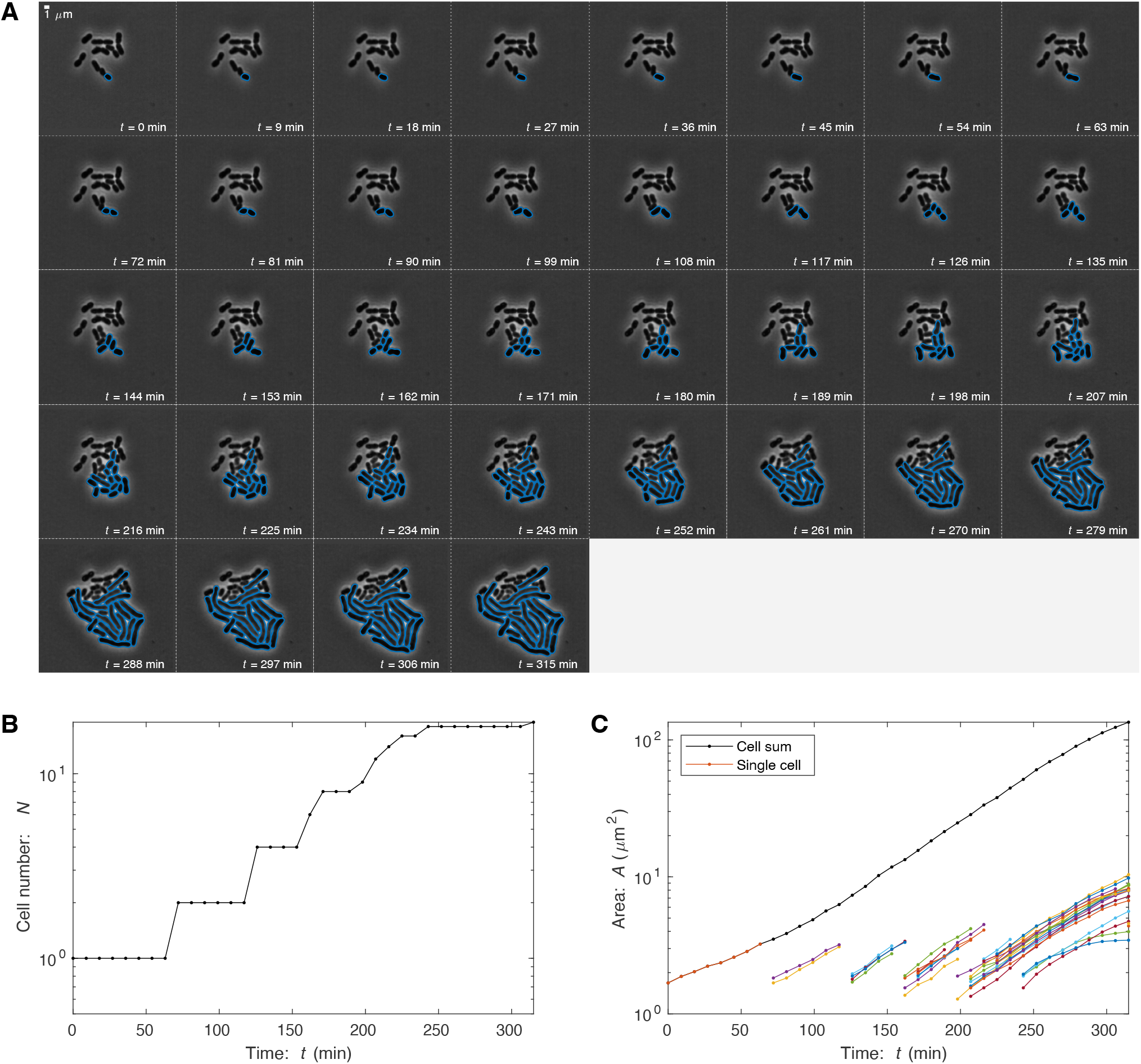
Knockout-depletion experiment: Δ*dnaN*. Panel A: Frame mosaic. One transformant (*dnaA*∷*kan*(Km^*R*^), blue) proliferates. The frame mosaic shows a typical imaging-based knockout-depletion experiment. DnaN is the sliding beta clamp, an essential DNA replication protein and a core component of the replisome. Its depletion leads to a failure of the chromosome to replicate and therefore results in cell filamentation. Cells were segmented using OmniSegger for quantitative analysis. **Panel B: Cell number**. The number of transformant progeny as a function of time. After transformation, normal growth persists for roughly 240 min, consistent with DnaN expression being overabundant. **Panel C: Progeny area**. Total progeny-cell area as a function of time. The areal elongation dynamics persists even are cell division arrests.

#### Supplemental approach

For this analysis, we did not want to explicitly model cell lysis. Therefore, in our fitting of the cell-number and areal growth curves, we locked the individual cell area at the last value taken immediately preceding lysis. Similarly, we treated cells that had lysed as arrested, not absent. (This fitting-refined data is *not* shown in Fig. S7. The resulting refined data for Panels B and C plateau rather than decease after growth arrest.)

**FIG S6.**
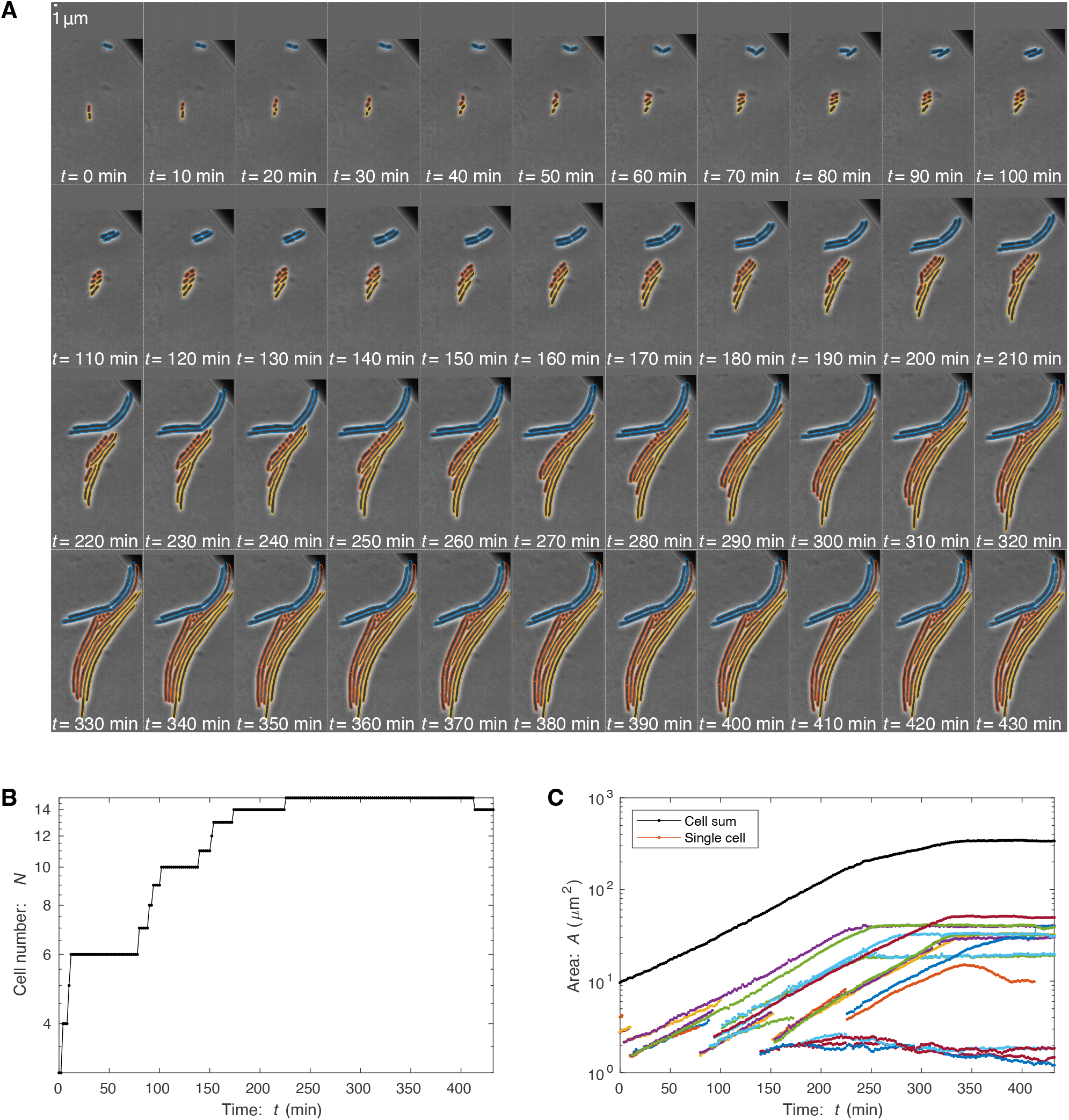
Knockout-depletion experiment: Δ*ftsN*. Panel A: Frame mosaic. Three transformants (*ftsN*∷*kan*(Km^*R*^), blue, yellow, orange) proliferate. FtsN is an essential cell division protein. Its depletion leads to a failure of the cells to septate. Cells were segmented using OmniSegger for quantitative analysis. **Panel B: Cell number**. The number of transformant progeny as a function of time. After transformation, normal growth persists for roughly 150 min, consistent with FtsN expression being overabundant. **Panel C: Progeny area**. Total progeny-cell area as a function of time. The areal elongation persists even after cell division arrests.

#### Quantitative analysis

The null hypothesis (*Sufficiency model*) was rejected for both the area (p-value under machine precision) and cell-number analysis (*p* = 1.2 ×10^−6^). The initial growth rate was observed to be *k* =0.70 ± 0.08 hr^−1^ with an arrest time of *T* = 3.6 ± 0.4 hr for the areal analysis. For cell-number analysis, the initial growth rate was observed to be *k* = 0.97 ± 0.24 hr^−1^ with an arrest time of *T* = 2.0 ± 0.3 hr.

## 2. *ACINETOBACTER BAYLYI* STRAINS, MANIPULATION, AND CULTURING

Mutant strains were derived from *Acinetobacter baylyi* ADP1 (MAY101) (the gift of C. Manoil) [41]. Growth media were LB and M9, a minimal-succinate M9 medium [42], supplemented with 15 mM sodium succinate, 2 mM magnesium sulfate, 0.1 mM calcium chloride and 1–3 µM ferrous sulfate (from sterile 5mM stock, made fresh at least once a month). For selective growth, media was supplemented with kanamycin at 20 µg/mL. Cultures were grown at 30°C.

The strains used in the study are summarized in Tab. S2.

### A. Methods: Construction of deletion mutations

We generated deletion mutants by transformation of linear DNA fragments, constructed by PCR using extension overlap [17]. A homologous overlap of ~2 kb flanking target genes was created that either directly joined (for marker-free deletions) or flanked a kanamycin resistance cassette (for kan-selectable deletions). Unmarked deletions were in-frame. Kan deletions were constructed from the *kan* gene from plasmid pACYC177 [43], in an orientation matching the deleted gene [17]. PCR reactions were performed using Q5 Polymerase (New England Biolabs) or Phusion HF polymerase (New England Biolabs) and DNA fragments were purified using Qiaquick columns (Qiagen) before transformation.

### B. Methods: *A. baylyi* transformation protocol

DNA fragments were transformed into *A. baylyi* cultures prepared as follows. Cultures were grown overnight in minimal-succinate M9 media with 1 µM ferrous sulfate. The culture was then back diluted 1:5 into fresh medium and grown one hour, shaking at 30°C. The DNA fragment was added at 1 µg/mL, followed by incubation for 2.5 - 3 hours with shaking, and then plated on selective (for *kan*-deletion cassettes) or non-selective media (for marker-free cassettes). Marker-free deletion mutants were identified by screening single colonies by PCR using primers flanking targeted genes. Essential gene kan-marked deletion mutations were selected by plating on protective medium supplemented with 20 µg/mL kanamycin. All unmarked and the marked non-essential deletion mutations were verified by PCR. For essential gene deletions, 0.1–1% of the cells were transformed, forming microcolonies of cells carrying the deletion.

### C. Methods: Construction of YPet-*dnaN* fusion strain

In previous work in *Escherichia coli* and *Bacillus subtilis*, we visualized fluorescent fusions to the beta sliding clamp (*dnaN*) to study replication [19–21]. The DnaN protein imaging is a convenient tool for studying replication due to its relatively high abundance and the change in its localization, from diffuse (non-replicating cells) to punctate (replicating cells), which serves as a convenient reporter of activity.

To construct a fluorescent fusion to the *A. baylyi* DnaN protein with a high probability of success, we used the exact same fluorescent protein and linker to that which R. Reyes-Lamothe had used to construct the *E. coli* fusion used in our previous work [25]. In this approach, we inserted the YPet-linker cassette at the 5’ end of the gene. Since the transformation efficiency of *A. baylyi* is so high, we constructed a marker-free fusion. We screened colonies by both PCR and fluorescence localization. We then sequenced the mutant *YdnaN* strain to confirm that the desired construct was achieved. (We provide a supplemental file with the sequence.) Like the original *E. coli* strain, no growth phenotype is observed under experimental conditions.

## 3. GROWTH MODELS FOR KNOCKOUT-DEPLETION EXPERIMENTS

To quantitatively analyze growth in knockout-depletion experiments, we define three nested growth models: (i) *No-Effect*, (ii) *Sufficiency*, and (iii) *Overabundance* models. In our statistical analysis, we will initially treat the No-Effect model as the null hypothesis and the Sufficiency model as the alternative hypothesis. If the null hypothesis is rejected, we will then adopt the Sufficiency model as the null hypothesis and adopt the Over-abundance model as the alternative hypothesis.

### 1. No-Effect model

In the *No-effect model*, the mutant has no effect on the growth rate. The abundance in a log culture will therefore be:

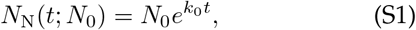

where *k*_0_ is the wild-type growth rate and *N*_0_ is the abundance at *t* = 0.

For modeling the TFNseq trajectories, it is the relative abundance that is measured and we therefore normalize by wild-type growth of the culture, resulting in the relative abundance:

**TABLE S2.**
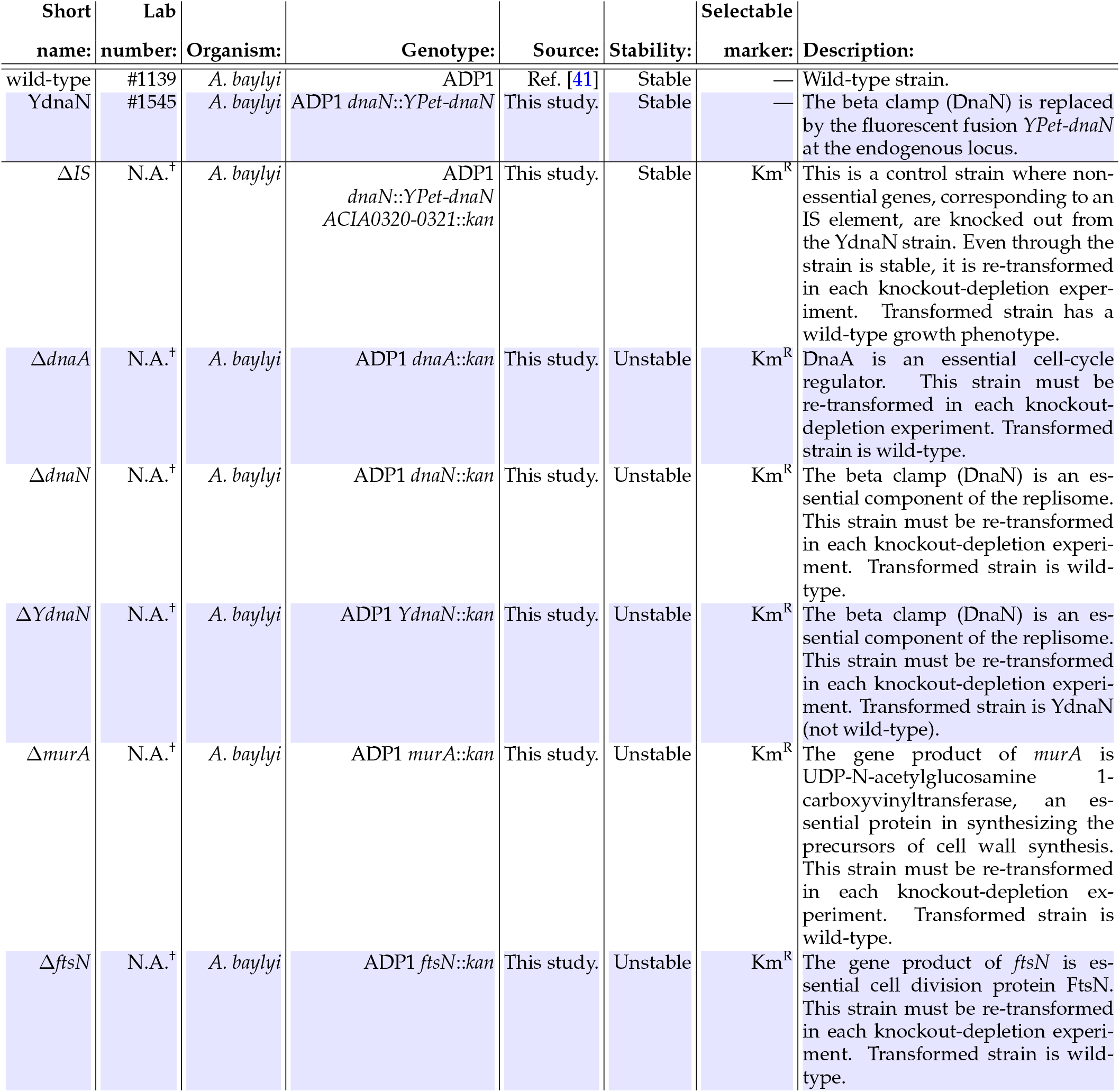
Summary of strains used in this study. The *short name* describes the nomenclature of the strains as described in the text. †Strain re-created by transformations in each knockout-depletion experiment are not stable and therefore are not assigned a *lab strain number* and, due to their instability, cannot be distributed.

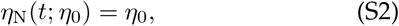

where *η*_0_ represents the initial relative abundance. (The relative abundance of the No-effect model is independent of *t*.) Both the abundance *N*_N_ and relative abundance *η*_N_ are plotted in Fig. S8. Both models depend on a single model parameter and are therefore dimension 1.

**FIG S7.**
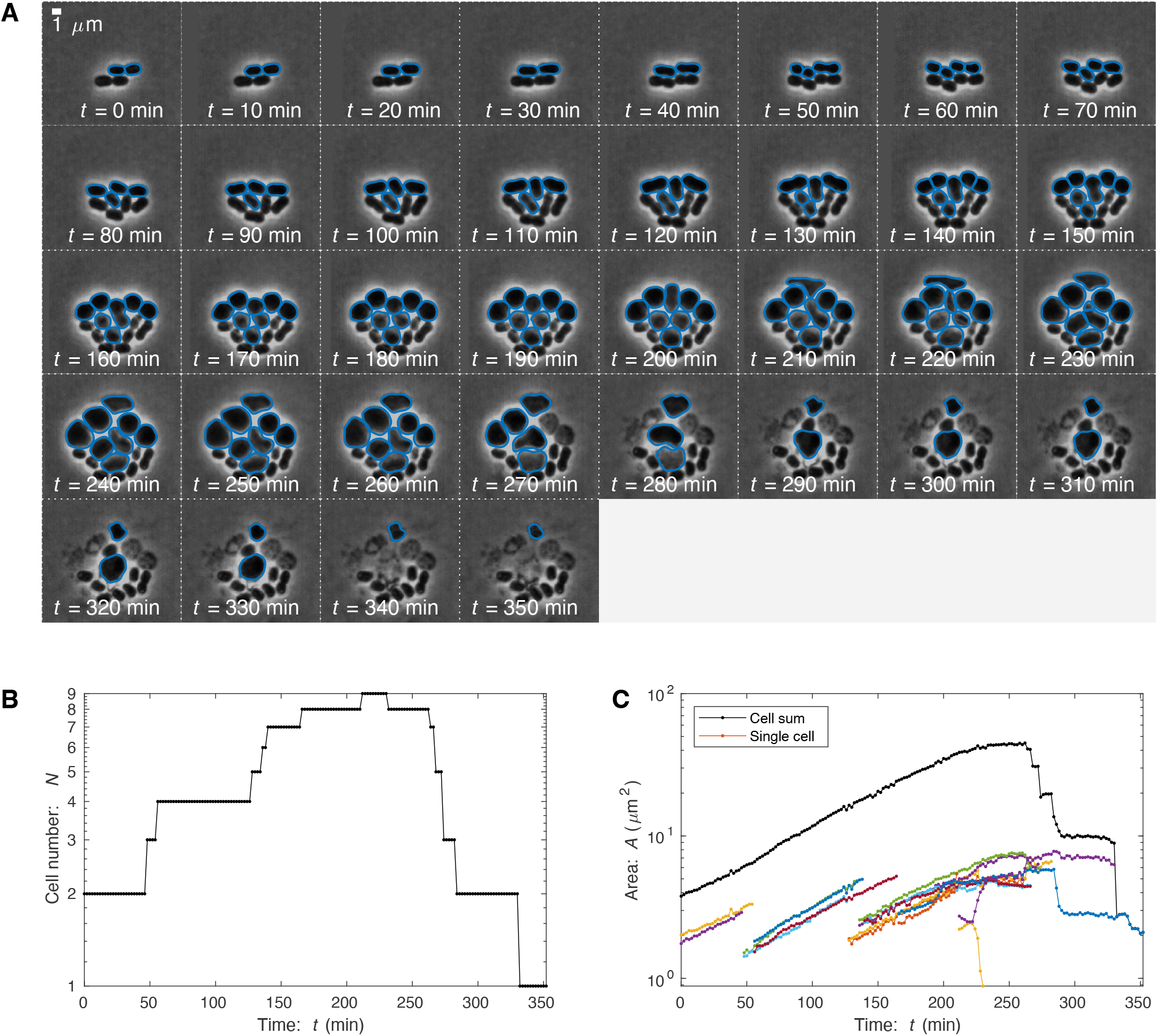
Knockout-depletion experiment: Δ*murA*. Panel A: Frame mosaic. Two transformants (*murA*∷*kan*(Km^*R*^), blue) proliferate. MurA is an essential enzyme responsible for cell wall precursor synthesis. Its depletion leads to the loss of cell wall integrity, and therefore first the loss of wild-type cell morphology and then cell lysis. Cells were segmented using OmniSegger for quantitative analysis. **Panel B: Cell number**. The number of transformant progeny as a function of time. After transformation, normal growth persists for roughly 200 min, consistent with MurA expression being overabundant. **Panel C: Progeny area**. Total progeny-cell area as a function of time. The areal elongation dynamics are largely consistent with the cell number dynamics: Normal growth persists for roughly 200 min.

### 2. Sufficiency model

In the *Sufficiency model*, we model the effect of the mutant as immediate. The cell number is assumed to grow at a new unknown rate:

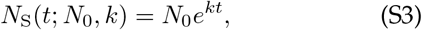

where *k* is the new growth rate and *N*_0_ is the number of the mutants at *t* = 0. For modeling the TFNseq trajectories, it is the relative abundance that is measured, and we therefore normalize by wild-type growth of the culture, resulting in the relative abundance:

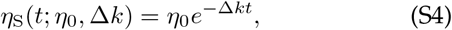

where Δ*k* ≡*k*_0_−*k* is the growth rate reduction of the mutant relative to the wild-type growth rate. Both the abundance *N*_S_ and relative abundance *η*_S_ are plotted in Fig. S8. Both models depend on two model parameters and are therefore dimension 2. Note that we might naïvely expect *k* = 0 for essential genes; however, we expect some transient growth due to residual protein levels, and these transients will dominate the fit.

**FIG S8.**
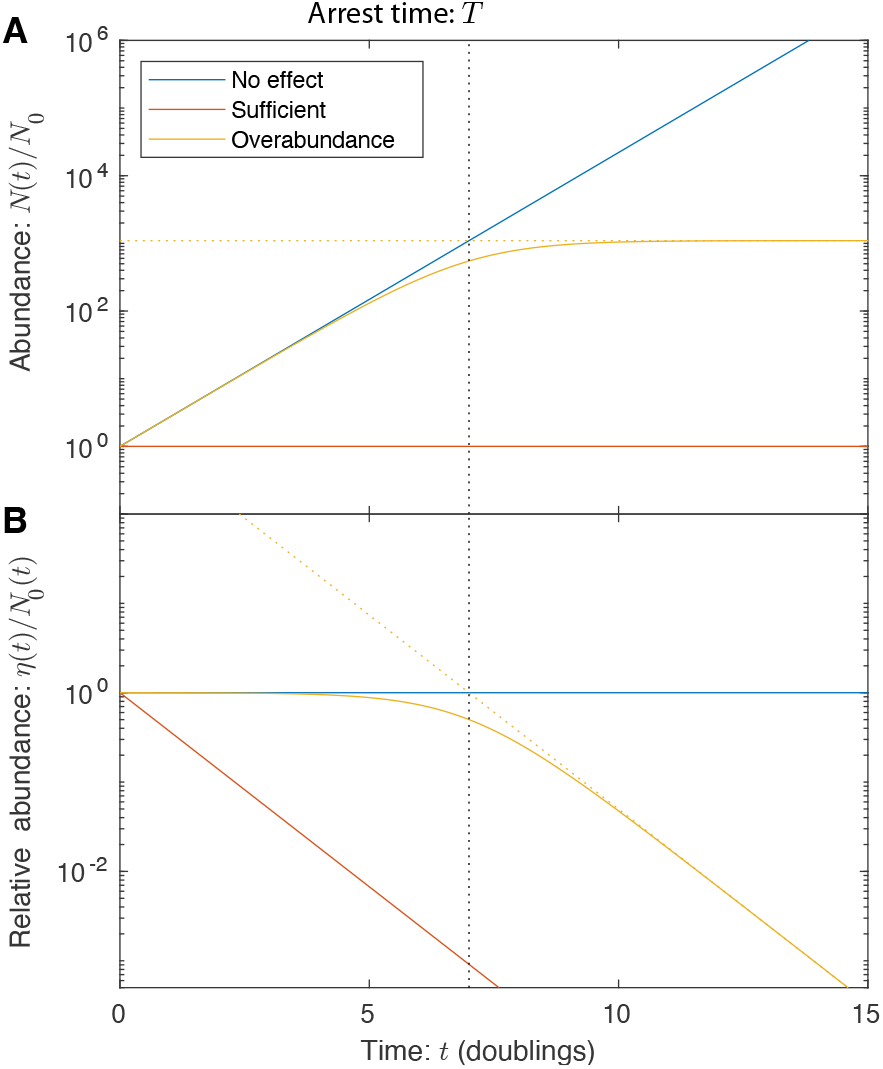
Panel A: Mutant abundances for trajectory models. Mutants described by the *No-effect model* (blue) grow at the wild-type growth rate. Mutants described by the *Sufficient trajectory model* (red) show an immediate change in growth rate after transformation. Mutants described by the *Overabundant trajectory model* (yellow) grow to the arrest time *T* (black dotted line) with the wild-type growth rate, before adopting a reduced growth rate of *k* = 0. **Panel B: Relative mutant abundances for trajectory models**. Same as above, but abundances are renormalized by wild-type growth.

### 3. Overabundance model

In the *Overabundance model*, we model the effect of the mutant with a delayed arrest time, *T*: the transient growth duration as protein dilutes to the threshold level. For short times, the mutant growth with a wildtype rate:

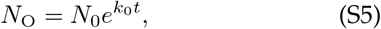

however, at long times we expect growth with a new unknown growth rate *k*:

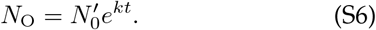

We initially attempted to use a piecewise function to join these two limits; however, the sparsity of the data and discontinuous slope at the boundary appeared to give rise to fitting artifacts. In addition, the cell-to-cell variation in protein expression smooths out the transition time. To fix these shortcomings, we adopted an empirical formula with the correct limits, but with a smooth transition at *t* = *T*:

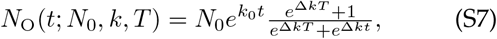

where Δ*k*≡ *k*_0_ −*k* is the loss in growth rate due to the mutation. Modeling the TFNseq trajectories, it is the relative abundance that is measured, and we therefore normalize by wild-type growth of the culture, resulting in the relative abundance:

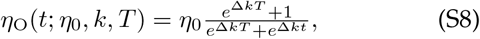

where Δ*k* ≡*k*_0_ −*k* is the growth rate reduction of the mutant relative to the wild-type growth rate. Both the abundance *N*_O_ and relative abundance *η*_O_ are plotted in Fig. S8. Both models depend on three model parameters and are therefore dimension 3. Note that we might naïvely expect *k* = 0 for essential genes; however, we expect some transient growth due to residual protein levels, and these transients will dominate the fit.

## 4. METHODS: IMAGING-BASED KNOCKOUT-DEPLETION EXPERIMENTS

### A. Methods: Experimental protocol

For single-cell imaging-based analyses, cells were imaged proliferating in M9 media supplemented with 2% low-melt agarose, and in most cases, kanamycin at 20 µg/mL.

#### 1. Cell preparation for knockout-depletion experiments

The transformation protocol described above was modified as follows: after the 2.5-3 hr incubation with DNA, cells were immediately spotted on selective media pads for imaging. In the knockout-depletion experiments, cells are transformed with knockout cassettes which recombine into the genome, resulting in Km^R^ knockout strains. If transformed cells are transferred to Km^+^ media too quickly, the competent cells do not have sufficient time to integrate the *kan* cassette before growth arrest. If cells are transferred too late, essential proteins are depleted before imaging begins. How do we know transformants after 2.5-3 hr outgrowth are at their initial stages of transient growth? With the 2.5-3 h outgrowth period, many cells still grow slowly (compared to log phase growth) for 10-15 min consistent with the expression of the kanamycin phosphotransferase (the gene product of the *kan* gene) not having reached a sufficiently high level to achieve a resistance phenotype. Furthermore, a significant number of heterogenic progenitors were observed. The presence of these heterogenic progenitor cells is consistent with the 2.5 h outgrowth period representing the typical recombination time for transformants. (See Sec. 1 A for a discussion of heterogenic progenitors.)

#### 2. Sample/slide preparation

Thin pads were fabricated by melting the agarose (Invitrogen UltraPure^TM^ LMP Agarose) and casting it between two slides with two layers of lab tape used as a shim to set the height. After the pad solidified (roughly 10 min), the top slide was carefully removed, and a razor blade was used to trim the pad to form a small square that could be covered with a #1.5 coverslip. For *E. coli* imaging, we typically use a pad that matches the size of the coverslip; however, *for A. baylyi imaging, we trim the pad so it is less than 1 cm in width. This added space allows aerobic growth to continue over multiple hours*. Finally, the coverslip is sealed using a hot glue gun.

#### 3. Microscopy

The samples were imaged using a Nikon Eclipse Ti microscope in phase contrast and fluorescence. We imaged through a Nikon 60 ×1.4 NA Phase contrast objective onto a sCMOS camera (Andor Neo). An environmental chamber maintained the sample at 30°C during imaging. For phase imaging, a frame rate of 1 frame / 2 min was used; however, for combined phase and fluorescence imaging we reduced the frame rate to 1 frame/ 3 min and 1 frame / 9 min to help reduce bleaching and phototoxicity. (The slowest frame rate was used to resolve the dim YPet-DnaN foci as the protein levels were depleted.) Typically, multiple (~10) fields of view were captured simultaneously in each experiment. For fluorescence-based analysis, we mixed in wild-type cells, in addition to fluorescent-fusion cells (1:2), to determine the autofluorescence levels in each experiment.

#### 4. Image processing (cell segmentation) pipeline

Cell images were processed using the *OmniSegger* package [44] by running the `processExp`command with default settings. Most of the analysis described in the paper was performed from the `clist.mat`files generated for each dataset.

### B. Methods: Cytometry data analyses

Imaging-based analysis for protein overabundance was carried out by assessing the transient cell area growth and septation. The three different single-cell analysis approaches are explained below: *protein abundance, area*, and *number analysis*.

#### 1. Accessing imaging-based cell cytometry data

Most of the analysis described in the paper was performed using the `clist.mat`files generated for each dataset by the *SuperSegger-Omnipose* package. In particular, the `data3D`field provides time-dependent cell descriptors for each cell in each frame, including `Rod Length, Area, Fluor1 sum`, and `Fluor1 background`. These descriptors were the input for our analyses. To characterize cell progeny area of fluorescence, we would generate cell lineage trees and cell progeny IDs using the `getFamily`command and then sum fluorescence or area over all progeny as a function of time. For instance, this data is shown in Fig. S3.

**FIG S9.**
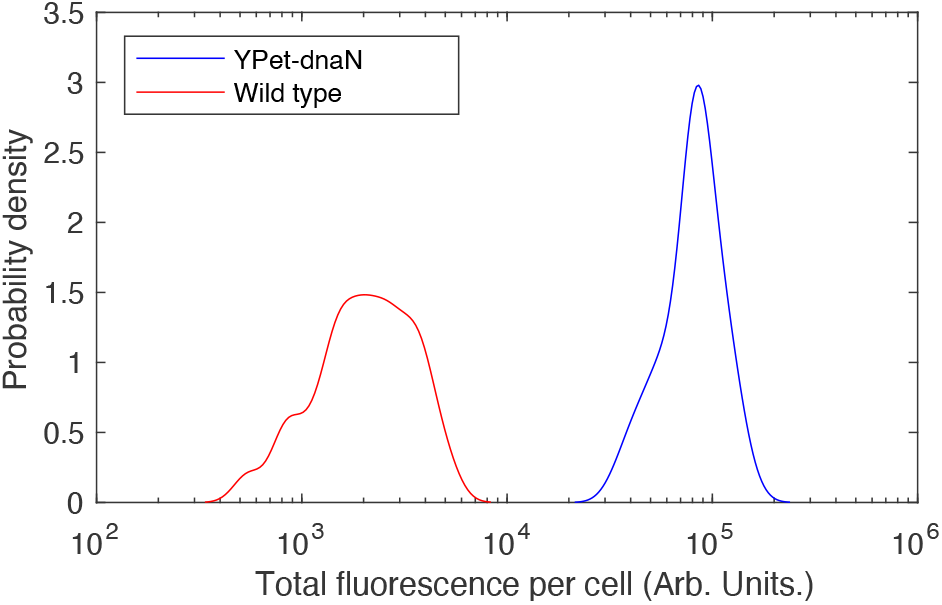
Wild-type and *YPet-dnaN* cell can be differentiated by fluorescence intensity with single cell resolution. Above we characterized fluorescence intensity for wild-type versus *YPet-dnaN* cells. Since the distributions are non-overlapping, the cells can be differentiated by fluorescence microscopy.

#### 2. Protein abundance analysis

To test the hypothesis that the targeted protein is depleted while protein-associated function continues for multiple generations, we visualized YPet-DnaN abundance and localization after the protein was knocked out as described in the paper. In short, we constructed a fluorescent fusion at the endogenous locus to make the YdnaN strain (Sec. 2 C), in which the endogenous *dnaN* was replaced by the fusion gene *YPet-dnaN*. In the knockout-depletion experiment, we knocked out the *YPet-dnaN* gene with the *kan* cassette to form *YPet-dnaN*∷*kan*.

We determined that wild-type and *YPet-dnaN* cells were unabiguously differentiable by fluorescence intensity. (See Fig. S9.)

To test the protein dilution hypothesis, we measured total progeny fluorescence (the proxy for protein abundance of YPet-DnaN) as a function of time, as the cell progeny proliferated. The dilution model predicts that the protein abundance should scale with the total progeny area like:

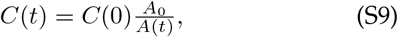

where *C*(*t*) is the protein concentration at time *t, C*_0_ is the abundance at time *t* = 0, *A*_0_ is the progenitor area at time *t* = 0, and *A*(*t*) is the total area of the progeny at time *t*. In the context of the fusion experiments, the observable is fusion fluorescence, equivalent to an intensity scaling of:

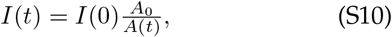

where *I*(*t*) and *I*(0) are the average pixel intensity of the progeny at time *t* and the progenitor at *t* = 0. Both area *A* and intensity *I* are time-dependent quantities available in the `clist.mat` file. (See Sec. 4 B 1.)

Several successive improvements in the experimental design and analysis were required to test the dilution hypothesis. (i) We initially attempted to image cells at the same frame rate as our phase contrast experiment (1 frame/2 min); however, to resolve YPet-DnaN foci after protein depletion, we had to significantly increase the exposure time of the fluorescence images and decrease the frame rate to avoid phototoxicity and bleaching. Although the predicted scaling (Eq. S10) was immediately observable in the data without corrections at short times, more care was required to observe the depletion at long times. (ii) First, we background subtracted to account for the background fluorescence level, computed as the average intensity in each frame outside the cell masks. This correction significantly improved the agreement with Eq. S10 at intermediate times, but did not yet account for cellular autofluorescence. (iii) Next, we analyzed a mixture of wild-type and *YdnaN* cells, using the intensity of the wild-type cell in the same microcolony for the background subtraction. This method led to good agreement with Eq. S10 even at long times (Fig. 1).

Why was a mixture of wild-type and *YdnaN* cells preferable to imaging the two strains independently? A detailed analysis of single cell intensities revealed that wild-type cells in close proximity to *YdnaN* cells in the microcolony had higher pixel intensity, due to the diffuse halo created by the bright *YdnaN* cells. The use of wild-type cells in the same field of view helped correct for the diffuse fluorescent light necessary for the analysis of protein abundance at large depletion times. Cell fluorescence intensities at *t* = 0 are used to differentiate between wild-type and *YdnaN* cells.

#### 3. Areal growth analysis

In this section, we develop the statistical model for the analysis of cell-area based growth assays to determine both the model parameters and the statistical uncertainty of parameters on a per-experiment basis. We provide this development for completeness; **however, cell-to-cell variation will dominate the reported errors**.

##### Statistical procedure

For the imaging-based analyses, we define the following statistical procedure: For the analysis of essential genes, we will fix the asymptotic growth rate *k* = 0. Therefore, the Sufficiency model is now considered the null hypothesis since it is the lowest dimensional model. The first alternative hypothesis is the No-effect model, where the wild-type growth rate *k*_0_ is fit in each analysis. If the Sufficiency model is rejected, we then adopt the No-effect model as the new null hypothesis and adopt the Overabundance model as the new alternative hypothesis.

##### Areal growth models

This growth metric is sensitive to cell elongation (rather than septation). Let *A*(*t*) be the observed area of all cells sharing a single progenitor cell. For the areal growth model, we substitute cell area *A*(*t*) for the abundance *N* (*t*) and *A*_0_ for *N*_0_ in Eqs. S1, S3, and S7. The models are:

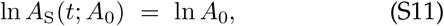

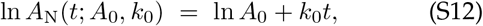

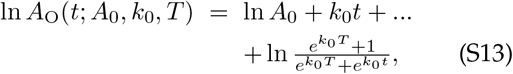

where we have substituted *k* = 0.

##### Statistical model for areal growth analysis

We will model the error associated with determining the area of the cells as proportional to cell number or area:

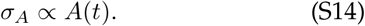

This model is consistent with many mechanisms. Rather than fitting a model with a variable error, it is more convenient to introduce a new variable, *a*, with constant error:

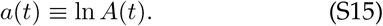

Since d*a* = d*A/A*, then *σ*_*a*_ = *σ*_*A*_*/A* which leads to an analysis with constant error.

The Shannon information (minus log likelihood) for the log area in frame *i* is:

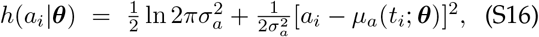

where ***θ*** represents the parameter vector, *µ*_*a*_ is the timedependent mean log area defined by the growth models (Eqs. S11-S13). For a time series with *i* = 1…*N* frames, the total Shannon information is:

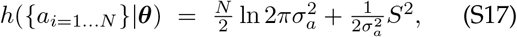

which can be formulated as a least-squares minimization where:

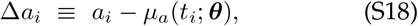

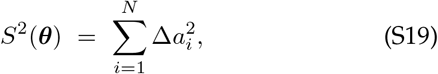

where *i* is the frame index.

##### Estimate of error for areal growth analysis

We will statistically estimate the relative area uncertainty (*σ*_*a*_) from the wild-type growth data. The expression for the MLE for 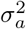 is:

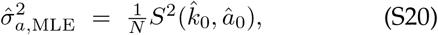

where Eq. S19 is evaluated at the MLE values of the other parameters for the No-effect model. There is one additional improvement to this estimate which is straight forward to implement. It is well known that Eq. S20 is biased from below. We can construct an unbiased estimator by correcting for the complexity of the model for the mean (dimension two) [28]:

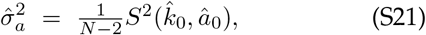

which we will use for our variance estimator. Note that if only a single mean were fit, the prefactor would be (*N* −1)^−1^ accounting for the one model dimension; however, since we fit both the slope and the offset, the prefactor is (*N*− 2)^−1^ accounting for the two model dimensions [28, 45].

From the wild-type growth data, the unbiased estimator for the error for log area (Eq. S21) is:

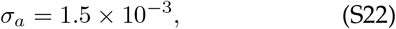

or alternatively, this result can be stated in a more intuitive form: There is a 0.15% error in the cell area.

##### Application to observed data

To determine the model parameters, we will minimize the Shannon information (Eq. S34) numerically, by a least-squares minimization of Eq. S18. We estimate the Fisher information using the resulting Jacobian from the least-squares minimization:

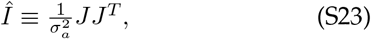

where the Jacobian matrices *J* are contracted over the frame index and *σ*_*a*_ is given by Eq. S22. The parameter uncertainties are then estimated from the Fisher information (Eq. S76). Although Eq. S76 accounts for the statistical uncertainty in the parameters, it does not account for the cell-to-cell variation. We found that this cell-to-cell variation was dominant. We therefore cite this cell-to-cell variation-based uncertainty. For the p-value calculations (Eq. S79), we compute the test statistic *λ* (Eq. S77) from the differences between residual norms for the null and alternative hypotheses:

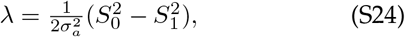

where *σ*_*a*_ is given by Eq. S22, and the residual norms for model I (the null (0) or the alternative (1) hypotheses) are defined in Eq. S19.

### C. Methods: Cell-number growth analysis

In this section, we develop the statistical model for the analysis of cell-number based growth assays to determine both the model parameters and the statistical uncertainty of parameters on a per-experiment basis. We provide this development for completeness; **however, cell-to-cell variation will dominate the reported errors**.

#### Statistical procedure

For the imaging-based analyses, we define the following statistical procedure: For the analysis of essential genes, we will fix the asymptotic growth rate *k* = 0. Therefore, the Sufficiency model is now considered the null hypothesis since it is the lowest dimensional model. The first alternative hypothesis is the No-effect model where the wild-type growth rate *k*_0_ is fit in each analysis. If the Sufficiency model is rejected, we then adopt the No-effect model as the new null hypothesis and adopt the Overabundance model as the new alternative hypothesis.

#### Cell-number growth models

For the cell-number growth model, we use Eqs. S1, S3, and S7. The statistical models depend on the growth rates as function of time for model *I*, which we define as:

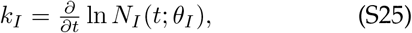

where *N*_*I*_ is the cell abundance in model *I* at time *t*. The growth rates for the respective models are:

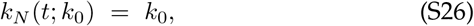

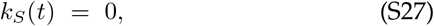

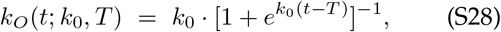

where Eq. S28 interpolates between the initial growth rate *k*_0_ and final growth rate *k* = 0 at time *T*.

#### Deriving the Shannon information

Consider an experiment in which images are taken with a high frame rate, where the time duration between frames is *δt*. Let the frame number be denoted *I* = 1…*m* and the number of cells in each frame *N*_*I*_. Let the model for cell growth be formulated such that the growth rate at time *t*_*I*_ is:

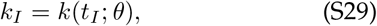

where *θ* represents a parameter vector. In this analysis, we with model cell division as a Markovian process where:

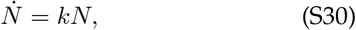

which is to say that we will ignore the internal state of cells. For instance, at time *t*, cells have the same rate of division, irrespective of cell age.

In this model, the number of cell divisions *n*_*I*_ that occur over the short time interval *δt* is Poisson distributed:

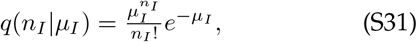

Where

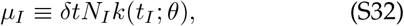

is the mean number of divisions.

We now compute the Shannon information associated with the entire experiment:

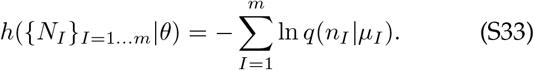

Substituting Eqs.S31 and S32, the equation is simplified to:

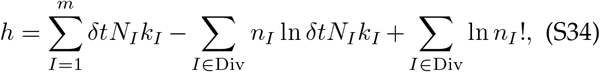

where Div represents the frames immediately preceding division. For instance, if there is one cell at frame 5 and two cells at frame 6, Div = {5}.

#### Application to observed data

To determine the model parameters (Eq. S75), we will minimize the Shannon information (Eq. S34) numerically, and determine the Hessian at the optimal parameter values to estimate the Fisher information:

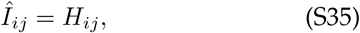

where *H* is the Hessian matrix. The parameter uncertainties are then estimated from the Fisher information (Eq. S76). We therefore cite this cell-to-cell variation-based uncertainty. For the p-value calculations (Eq. S79), we compute the test statistic *λ* (Eq. S77) from the differences in the Shannon information (Eq. S34).

#### Error estimates of arrest times in imaging-based approach

Due to the noisiness of gene expression, there is significant cell-to-cell variation in protein abundance from progenitor to progenitor in the image-based analysis; however, there is significantly less noise in protein partitioning between the daughter cells [22]. Eq. S76 accounts for the statistical uncertainty in the parameters for a single progenitor cell, but it does not capture the uncertainty due to cell-to-cell variation. To estimate the uncertainty associated with cell-to-cell variation, we used the canonical unbiased empirical error estimator [28]:

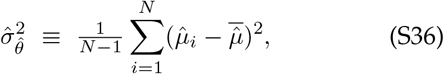

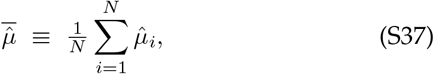

where index *i* runs over the progenitor cells 1..*N* and 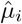 is the estimate of the parameters from progenitor *i*. We found that this cell-to-cell variation (Eq. S36) was larger than the per-progenitor estimated error (Eq. S76).

## 5. STATISTICAL ANALYSIS OF TFNSEQ TRAJECTORIES

### A. Methods: Time correction

Since the mutants transition from lag phase to log phase after transformation, we used a log phase equivalent time for the TFNseq-approach analysis. The corrected sampling times (*t*_*s*_) are estimated from the number of doublings (*D*_*s*_) for the non-essential mutants obtained from TFNseq experiment([18]):

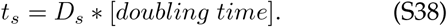

For our experiment, the doubling time for ADP1 in M9 at 30°C is 37 min.

### B. Methods: Defining the likelihood

We assume that deep-sequencing is well modeled by a Poisson process for which the probability mass function is:

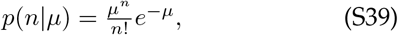

where *n* is the number of reads and *µ* is the mean-number parameter. For large *n*, we use the normaldistribution approximation:

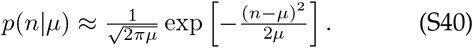

The total likelihood for sequential observations *n*_1…*m*_ at time *t*_1…*m*_ is therefore:

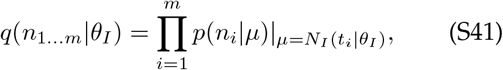

where *N*_*I*_ is one of the trajectory models and *θ*_*I*_ is the parameter vector for model *I*. The Shannon information is:

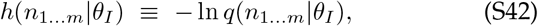

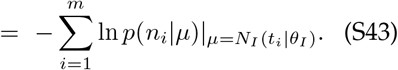

To fit this expression, we use the total number of reads mapped to each mutant gene as the *n*_*i*_ and minimize to determine the MLE parameter estimates.

### C. Methods: Data binning for overabundance plot

We used two different binning approaches to show the experimental trend in overabundance (*o*) as function of transcription (*µ*_*m*_).

#### 1. µ-binning

For the *µ*-binning approach, we used the canonical approach: We divided the log-10 interval of *µ*_*m*_ into bins width Δ log_10_ *µ*_*m*_ = 0.5. The resulting binned is shown in Fig. S12A.

The short-coming with this approach is that there is uncertainty in both *x* and *y* axes (transcription measurements (message number *µ*_*m*_) and overabundance (*o*) respectively) and there is a steep slope of the predicted message number as *µ*_*m*_→ 1 which obscures the trend for the data at low transcription levels.

**FIG S10.**
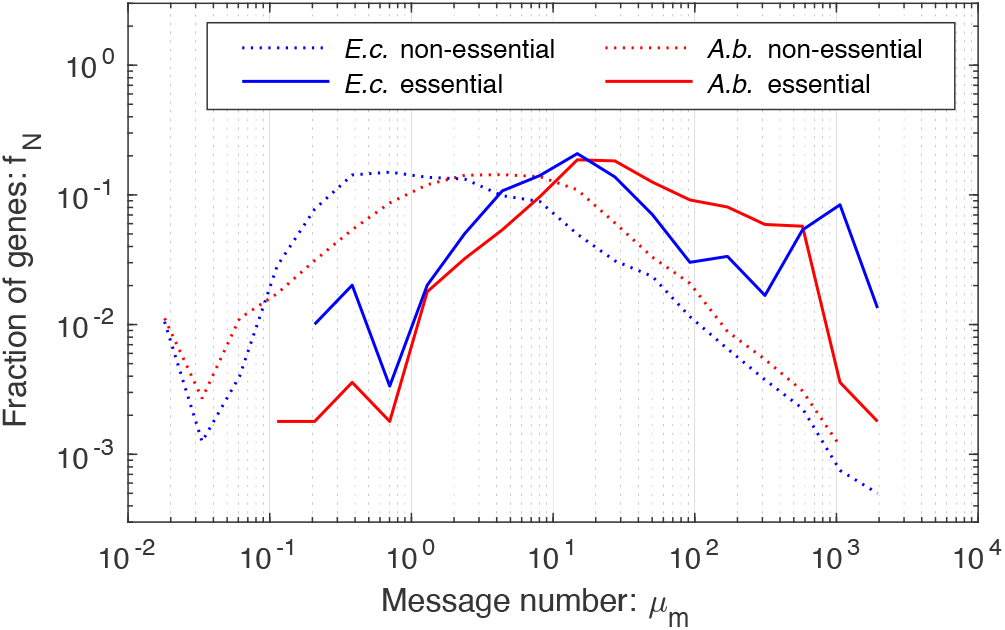
Message number distributions for essential and non-essential genes in *E. coli* and *A. baylyi*. Nearly all *A. baylyi* essential genes are expressed above the one-message-per-cell-cycle threshold. This distribution of both non-essential and essential genes in *A. baylyi* is qualitatively similar to that in *E. coli*, as predicted [11].

#### 2. n-binning

To better capture the data trend as *µ*_*m*_→ 1, we binned the data using an alternative dependent variable, *n*_*m*_. The threshold transcription level *n*_*m*_ is defined as:

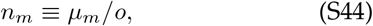

and can be understood as the transcription level that would be required in RLTO model, in the absence of gene-expression noise, for growth. We used an identical for *n*_*m*_: Δ log_10_ *n*_*m*_ = 0.5. The resulting binned is shown in Fig. S12B.

### D. Results: Overabundance is consistent between two knockout-depletion replicate experiments

Two independent knockout-depletion experiments were performed [18]. Qualitatively, the data from Experiment 1 fit the kinetic model better and therefore we used this dataset our primary analysis. However, the entire analysis was repeated for Experiment 2 data and the results are qualitatively unchanged. The overabundances for both experiments are reported in the Supplementary Data S1. The inferred repilcate overabundance values are compared in Fig. S13.

We used the second dataset to estimate experimental uncertainties in the reported log overabundance values by estimating the variance from the two observed data points for each gene *i*:

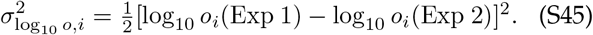

To quantify the typical experiment-to-experiment variation, we computed the gene-averaged variance in the log overabundance over essential genes. The variance and standard deviation are:

**TABLE S3.** Gene ontology classifications and terms. A summary of the gene ontology classifications and terms used in the study.

| Class | Terms |
| --- | --- |
| GO:0006260 | DNA replication |
| GO:0051301 | Cell division |
| GO:0008610 | Lipid biosynthetic process |
| GO:0009252 | Peptidoglycan biosynthetic process |
| GO:0008643 | Carbohydrate transport |
| GO:0006355 | Regulation of DNA-templated transcription |
| GO:0003824 | Catalytic activity |
| GO:0003677 | DNA binding |

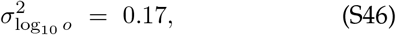

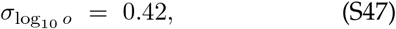

which corresponds to a multiplicative uncertainty of roughly 2.6-fold in the overabundance. We conclude that the inferred overabundance are broadly consistent between the experiments.

### E. Methods: Detection limit

We define the detection limit *λ*_min_ for the TFNseq experiments as the Poisson rate at which the mutant would be detected 95% of the time. This corresponds to the probability of non-detection (i.e. not measuring a count) as 5%. Therefore:

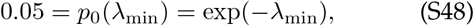

where *p*_0_ is the Poisson probability for zero counts. The detection limit is therefore:

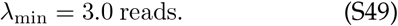

### F. Methods: Analysis of overabundance for different Gene Ontologies (GO)

To classify genes, the gene ontology classifications and terms summarized in Tab. S3 were used.

We analyzed only genes in *A. baylyi* that had homologues in *E. coli*. The *E. coli* classifications were downloaded from EcoCyc database [46].

### G. Analysis of overabundance for different gene regulatory controls

To investigate the effect of transcriptional regulation in determining protein overabundance, we assumed that the regulatory network in *A. baylyi* is roughly equivalent to that in *E. coli* which has been much more extensively studied. We used the EcoCyc database [46] to generate a list for each gene *i* of the list of direct regulators. For each gene, we counted the direct regulators of each gene, then ranked the genes in terms of regulator number, and finally we defined the top 10% of the genes as *highly regulated*. We also generated a list of genes directly inhibited by each gene. If a gene directly inhibited itself, we defied the gene as *autoregulatory*.

**FIG S11.**
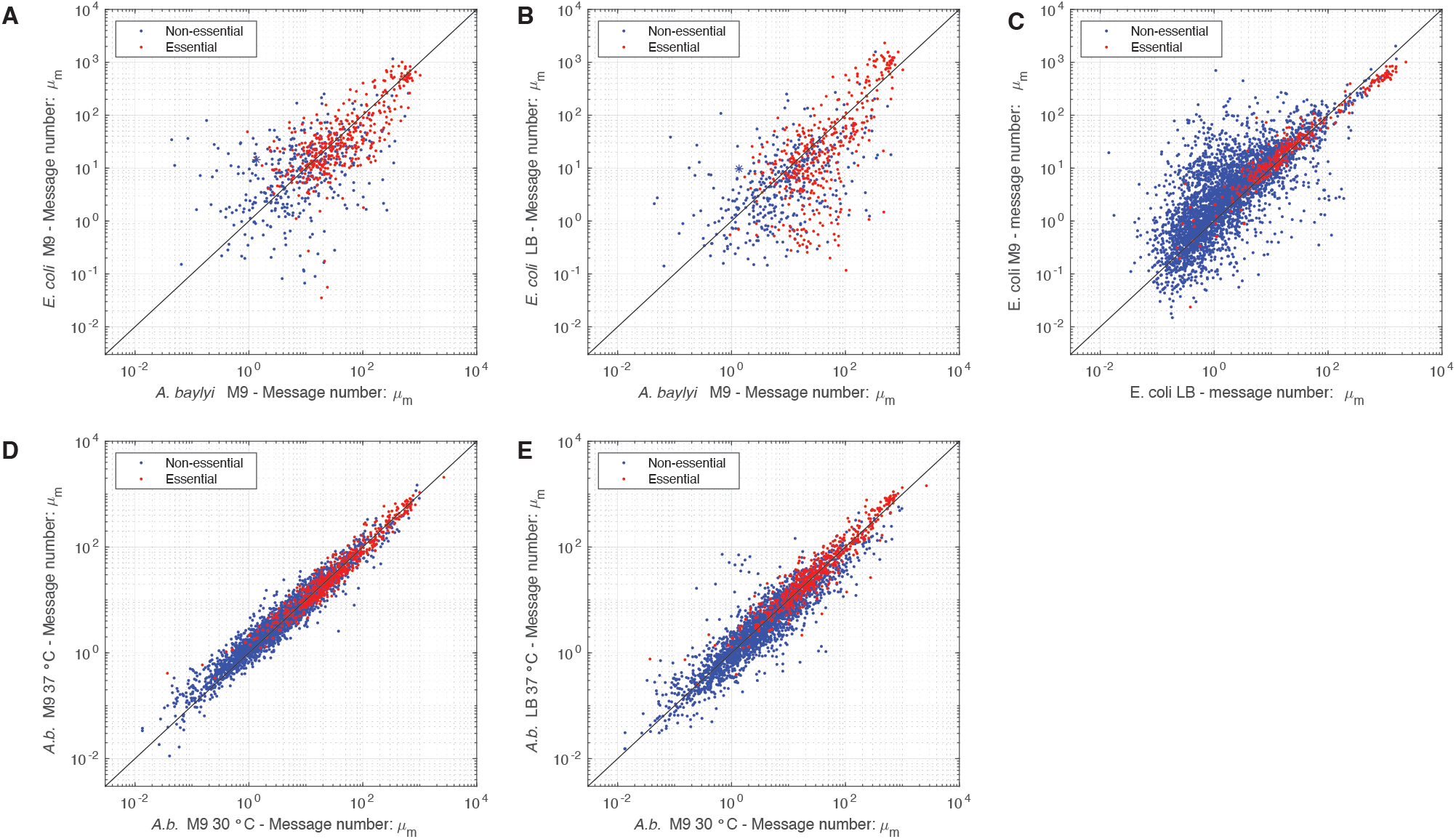
Transcriptome comparisons. Panel A: *E. coli* on M9 versus *A. baylyi* on M9. Panel B: *E. coli* on LB versus *A. baylyi* on M9. Panel C: *E. coli* on M9 versus on LB. Panel D: *A. baylyi* on M9 at 37°C versus at 30°C. Panel E: *A. baylyi* on LB at 37°C versus on M9 at 30°C. Throughout, there is broad consistency between the expression levels (message number) of genes, both between organisms and between conditions. These observations suggest a consistent overall transcriptional program governs gene expression both between organisms and growth conditions.

## 6. RNA-SEQ ANALYSIS OF TRANSCRIPTION

### A. Methods: RNA-Seq protocol

#### 1. RNA extraction

ADP1 RNA was harvested through methods developed by Culviner et al. [47] Total RNA was harvested by mixing 1ml of *A. baylyi*( 0.5 OD) with 110ul of icecold stop solution (95% ethanol and 5% acid-buffered phenol) and spinning in a tabletop centrifuge for 30 s at 13,000 rpm. The supernatant was flash-frozen and stored at −80°C until RNA extraction is ready. To start RNA extraction, 1ml of heated 65°C was added to the sample. The mixture was shaken at 65°C for 10 min and flash-frozen at −80°C for at least 10 min. The pellets were thawed at room temperature and spun at top speed in a benchtop centrifuge at 4°C for 5 min. The supernatant was collected and added to 400 µl of 100% ethanol. The mixture was passed through DirectZol spin column (Zymo). The column was washed twice with RNA prewash buffer and once with RNA wash buffer (Zymo). RNA was eluted from the column with 90 µl diethyl pyrocarbonate (DEPC)-H2O. Genomic DNA was removed with 4 µl of Turbo DNase I (Invitrogen) and supplemented with 10µl of 10x Turbo DNase I buffer to a final volume of 100µl. The solution was heated to 37°C for 40 min. Then RNA was diluted with 100 µl DEPC-H2O, extracted with 200 µl buffered acid phenolchloroform, followed by ethanol precipitation at −80°C for 4 h with 20 µl of 3 M sodium acetate (NaOAc), 2 µl GlycoBlue (Invitrogen), and 600 µl ice-cold ethanol. To pellet RNA, the samples were centrifuged at 4°C for 30 min at 21,000 × g. The pellets were washed twice with 500 µl of ice-cold 70% ethanol, followed by centrifugation at 4°C for 5 min. RNA pellets were then air dried and resuspended in 50 µl DEPC-H2O. The yield and integrity of RNA was verified with NanoDrop spectrophotometer, and by running 50 ng of total RNA on a Novex 6% Tris-buffered EDTA (TBE)-urea polyacrylamide gel (Invitrogen).

**FIG S12.**
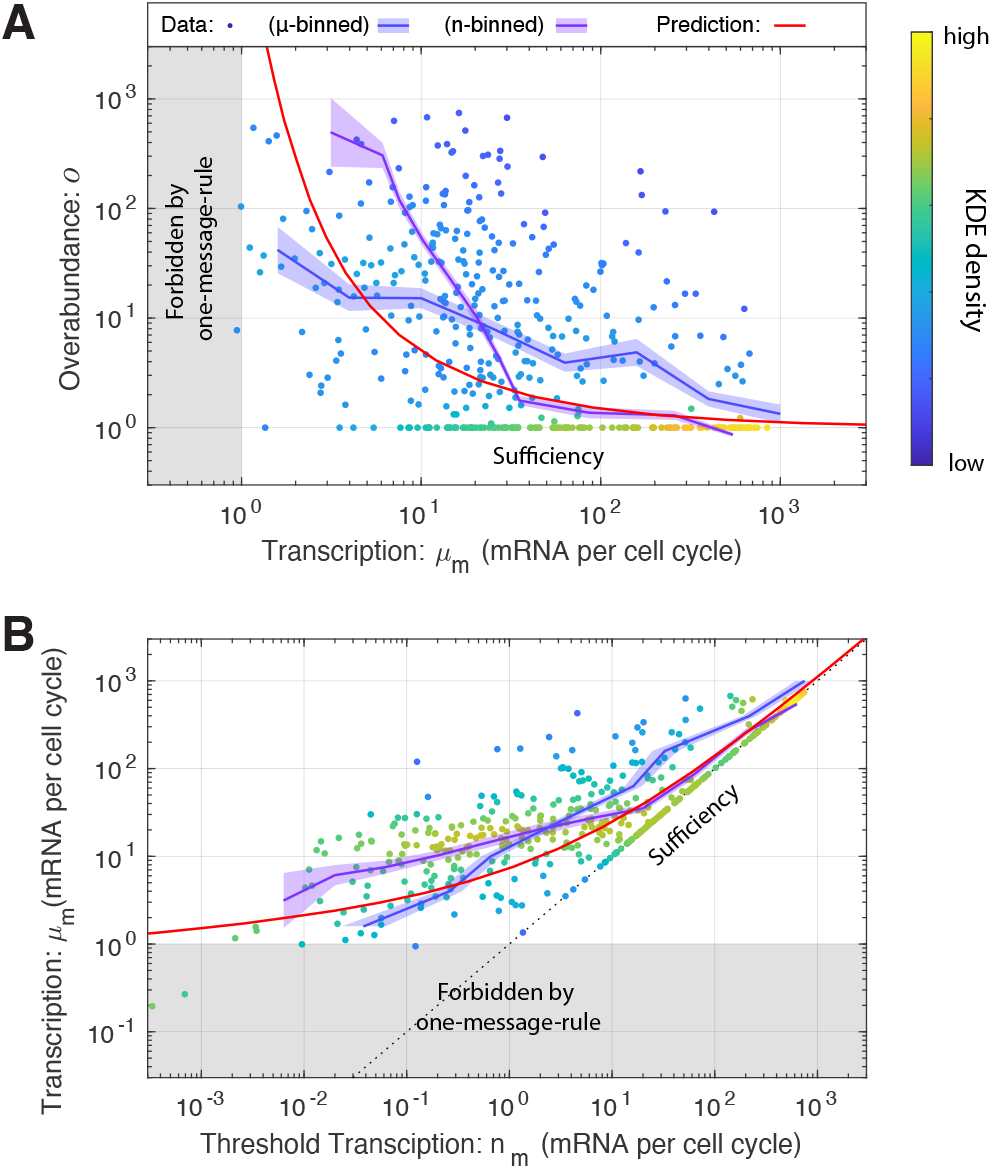
Panel A: Overabundance versus transcription. The measured transcription-overabundance pairs are shown for essential genes (including estimated gene density.) The blue curves shows the *µ*-binned data. The RLTO model (red) predicts that overabundance grows rapidly as the transcription level is reduced. **Panel B: Transcription versus threshold transcription**. Alternatively, the message number *µ*_*m*_ is shown as a function of the inferred threshold message number: *n*_*m*_ ≡ *µ*_*m*_*/o*. The purple curve shows the *n*-binned data.

**FIG S13.**
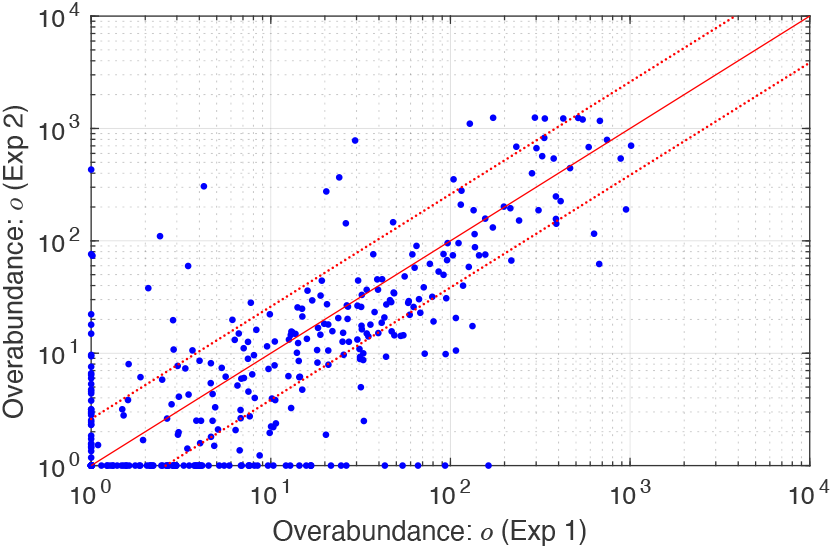
Overabundance is consistent between two knockout-depletion replicate experiments. The inferred overabundance values are compared for two experimental replicates. Perfectly matched experiments would align on the diagonal (red). The inferred overabundance values are broadly consistent between the two experiments.

#### 2. rRNA Depletion

rRNA was depleted through the DIY method developed by Culviner et al. [47] as well. We used their 21 biotinylated oligonucleotides for *E. coli*. The selected biotinylated oligonucleotides were synthesized by IDT and resuspended to 100 µM in TE buffer (Qiagen). An oligonucleotide mixture was made by mixing equal volumes of each 16S and 23S primers and double volumes of 5S primers. The pooled mixture was diluted with DEPC treated H2O based on the total RNA, using their Excel-based calculator. Using the Excel-based calculator, the calculated volume of Dynabeads MyOne streptavidin C1 beads (ThermoFisher) were washed three times in equal volume of 1x B&W buffer, resuspended in 30 µl of 2× B&W buffer and supplemented with 1µl of SUPERase-In RNase inhibitor (ThermoFisher). The beads were set aside in room temperature until the probes were ready to be pulled down. To collect rRNA, 2 to 3µg total RNA and 1 µl of the diluted biotinylated probe mix were combined on ice into a final annealing reaction mixture of 1xSSC and 500 µM EDTA. All the appropriate volumes were computed using the Excel-based calculator. The RNA and probe mixture was incubated at 70°C for 5 min, and slowly cooled to 25°C at a rate of 1°C per 30 s. The annealed mixture was then added to 30 µl of beads that were resuspended in 2× B&W buffer. The mixture was mixed by pipetting and vortexing at medium speed, and followed by incubating for 5 min at room temperature. The reaction mixtures were then vortexed, and incubated at 50°C for 5 min. To pull down the biotinylated probes, the reaction mixtures were placed immediately placed on the magnetic rack. The supernatant was carefully pipetted, placed on ice, and diluted to 200 µl in DEPC-H2O. The RNA was purified through ethanol precipitation with 20 µl of 3 M NaOAc, 2 µl GlycoBlue (Invitrogen), and 600 µl ice-cold ethanol at −20°C for at least 1 h. To pellet RNA, the samples were centrifuged at 4°C for 30 min at 21,000 × g. The pellets were washed twice with 500 µl of ice-cold 70% ethanol, followed by centrifugation at 4°C for 5 min. RNA pellets were then air dried and resuspended in 10 µl DEPC-H2O. The yield and rRNA depletion effectiveness was verified with NanoDrop spectrophotometer, and by running 50 ng of total RNA on a Novex 6% Tris-buffered EDTA (TBE)-urea polyacrylamide gel (Invitrogen). The yield and integrity of the library was checked by running the samples in qPCR using NEBNext Library Quant Kit for Illumina(NEB) and the Bioanalyzer.

#### 3. Library prep and sequencing

The RNA library was prepared with NEBNext® Multiplex Oligos for Illumina(NEB) and NEBNext ultra II RNA Library Prep Kit for Illumina(NEB). For the library prep protocol, we followed section 4 of the kit’s provided protocol: Protocols for use with Purified mRNA or rRNA Depleted RNA. The quality of the final library was verified by running the samples on high sensitivity Bioanalyzer chip. The samples were pooled to a final concentration of 8.5nM, and were sequenced with NextSeq 150 cycle kit.

### B. Methods: Computation of message number

To estimate the message number for gene *i*, defined as the total number of mRNA molecules transcribed per cell cycle, from the RNA-Seq data, we use the approach we described earlier [11]. Let the relative number of reads for gene *i* be *r*_*i*_:

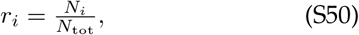

where *N*_*i*_ is the reads per kilobase (rpk) for gene *i* and *N*_tot_ is the rpk for all genes. We apply two different re-scalings: First we re-scale the relative message abundance to reflect the cellular abundance of the message, and then we scale this number by the ratio of cell cycle duration to mRNA lifetime to estimate the number of times a gene is transcribed per cell cycle. For *A. baylyi*, we use the same scaling factor as *E. coli*:

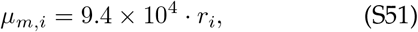

where *µ*_*m,i*_ is the estimated message number (number of mRNA molecules transcribed per cell cycle).

To check the consistency of this estimate, we generated histograms for message number for essential and non-essential genes, and compared them to the histograms for *E. coli*. We expect the distribution of essential message numbers to abut 1 message per cell cycle, while non-essential genes can be expressed at significantly lower levels. The observed distribution are consistent with this expectation. (See Fig. S10.)

### C. Results: Comparison of *A. baylyi* and *E. coli* gene expression

Knockout-depletion experiments are not tractable in *E. coli* and many other model systems. It is therefore difficult to directly test the overabundance hypothesis in these other systems. However, it is possible to determine if *E. coli* expression patterns are consistent with overabundance.

If overabundance were specific to *A. baylyi*, we would expect to see higher relative transcription of lower abundance essential genes in *A. baylyi*, where overabundance is large, relative to *E. coli* if its expression levels were sufficient. Fig. S11 compares the message number between homologues in the two organisms and between growth conditions within a particular organism for all genes.

## 7. ROBUSTNESS LOAD TRADE-OFF (RLTO) MODEL

We have provided a detailed description of the Robustness Load Trade-Off (RLTO) Model in Ref. [11]; however, in the interest of making this paper selfcontained, we provide a concise summary of key elements and results from that paper in this supplementary section.

### A. Methods: Detailed description of the noise model

#### 1. Stochastic kinetic model for the central dogma

The canonical steady-state noise model for the central dogma describes multiple steps in the gene expression process [34, 35, 48]: Transcription generates mRNA messages. These messages are then translated to synthesize the protein gene products [49]. Both mRNA and protein are subject to degradation and dilution [50]. At the single cell level, each of these processes are stochastic. We will model these processes with the stochastic kinetic scheme [49]:

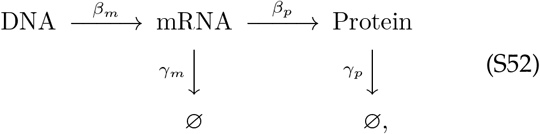

where *β*_*m*_ is the transcription rate (s^−1^), *β*_*p*_ is the translation rate (s^−1^), *γ*_*m*_ is the message degradation rate (s^−1^), and *γ*_*p*_ is the protein effective degradation rate (s^−1^). The message lifetime is 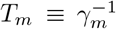. For most proteins in the context of rapid growth, dilution is the dominant mechanism of protein depletion and therefore *γ*_*p*_ is approximately the growth rate [48, 51, 52]: *γ*_*p*_ = *T* ^−1^ ln 2, where *T* is the doubling time.

#### 2. Statistical model for protein abundance

Consistent with previous reports [34, 35], we find that the distribution of protein number per cell (at cell birth) was described by a gamma distribution [11]:

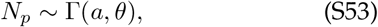

where *N*_*p*_ is the protein number at cell birth and Γ is the gamma distribution, which is parameterized by a scale parameter *θ* and a shape parameter *a*. (See Sec. 9 A.) We refer to this distribution as the *canonical steady-state noise model*; The relation between the four kinetic parameters and these two statistical parameters has already been reported, and have clear biological interpretations [35]: The scale parameter:

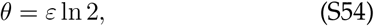

is proportional to the translation efficiency:

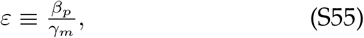

where *β*_*p*_ is the translation rate and *γ*_*m*_ is the message degradation rate. *ε* is understood as the mean number of proteins translated from each message transcribed. The shape parameter *a* can also be expressed in terms of the kinetic parameters [35]:

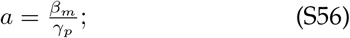

however, we will find it more convenient to express the scale parameter in terms of the cell-cycle message number:

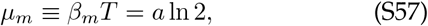

which can be interpreted as the mean number of messages transcribed per cell cycle. Forthwith, we will abbreviate this quantity *message number* in the interest of brevity.

### B. Methods: Summary of the RLTO model fitness model

#### 1. Metabolic load in the RLTO model

To produce a minimal model to study the trade-off between robustness and metabolic load, we must consider both the metabolic cost of transcription and translation. We will write that the metabolic load (in protein equivalents) associated with gene *i* is:

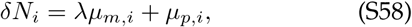

where *λ* is the message cost, the metabolic load associated with an mRNA molecule relative to a single protein molecule of the gene product.

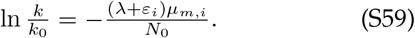

This equation has an intuitive interpretation: growth slows in proportion to the relative added metabolic load. In resource allocation models [53], the capacity of the cell for growth can increase as protein sectors increase in size. In our context, this does not occur since we consider the uncoordinated changes in the levels of single proteins. *I*.*e*. we assume some other protein of factor is rate limiting. See the detailed discussion in Ref. [11].

#### 2. Growth rate with stochastic arrest

As discussed in Ref. [11], we idealize the slow growth associated with essential proteins falling below threshold as growth arrest. This arrest model has phenomenology consistent with more detailed and realistic models where cells experience a significant growth slowdown rather than true growth arrest [11].

In the idealized growth arrest model, if all essential proteins are above threshold, the cell cycle duration *τ* is determined by the metabolic load predictions (Eq. S59); however, if any essential protein is below threshold, the cell cycle duration is infinite. The probability mass function for the cycle-cycle duration *T* interpreted as a random variable is therefore:

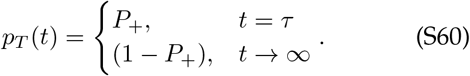

As we show in Ref. [11], the growth rate can be computed exactly:

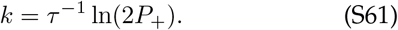

As expected, the growth rate goes down as the probability of growth *P*_+_ decreases, stopping completely at 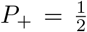. We can then compute the ratio of the growth with (*k*) and without arrest (*k*_0_):

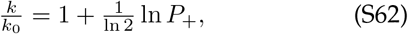

where *k*_0_ is computed by evaluating Eq. S61 at *P*_+_ = 1.

#### 3. RLTO growth rate

In the RLTO model, we will assume the probability of growth is the probability that all essential protein numbers are above threshold. We will further assume that each protein number is independent, and therefore:

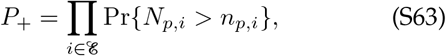

where ℰ is the set of essential genes. Clearly, this assumption of independence fails in the context of polycistronic messages. We will discuss the significance of this feature of bacterial cells elsewhere, but we will ignore it in the current context. As we will discuss, the probability of arrest of any protein *i* to be above threshold is extremely small. It is therefore convenient to work in terms of the CDFs, which are very close to zero:

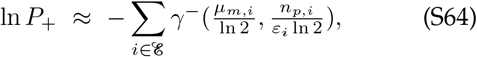

where *γ*^−^ is the regularized lower incomplete gamma function (Eq. S85) and represents the probability of arrest.

#### 4. Single-gene equation

By summing the fitness losses from the metabolic load and cell arrest (Eqs. S59, S62, and S64), we can write an expression for the growth rate including contributions from essential gene *i*:

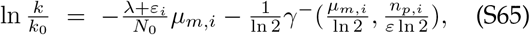

where the first term on the RHS represents the fitness loss due to the metabolic load and the second term represents the fitness loss due to stochastic cell arrest due to protein *i* falling below threshold.

#### 5. Optimization of transcription for bacteria

The growth rate is:

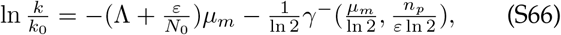

where *γ*^−^ is the regularized lower incomplete gamma function (Eq. S85), which is the CDF of the gamma distribution and represents the probability of arrest due to gene *i*. For bacteria, we consider the special case of optimizing the message number only at fixed translation efficiency [11, 48]. To determine the optimal transcription level, we set the partial derivative of Eq. S66 with respect to *µ*_*m*_ to zero. The optimum message number 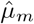 satisfies the equation:

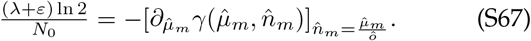

We define the relative load:

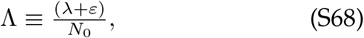

and substitute this into the optimum message number equation:

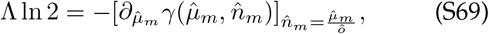

which is solved numerically.

#### 6. Estimate of the relative load in bacterial cells

In bacterial cells, we will assume a constant translation efficiency model. We therefore use the modified relative load formula (Eq. S68) to estimate Λ. We will assume that the load is dominated by proteins and messages:

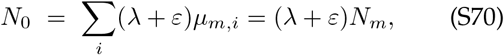

where *N*_*m*_ is the total number of messages. We can then solve this equation for Λ:

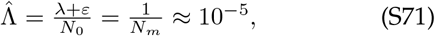

based on the total message number estimate for *E. coli* [11].

### C. Results: The fitness landscape of the RLTO model is highly asymmetric

In the RLTO model, the fitness landscape for a single cell is determined by an asymmetric fitness landscape: protein underabundance is extremely costly due to the risk of growth arrest, while the cost of protein overabundance is only associated with an increase in metabolic load. (See Fig. S14A.) Naïvely, this tradeoff predicts that the cell maximizes its fitness by simply expressing just above the minimum protein threshold for function [7]. However, achieving growth robustness at a population level is nontrivial. Gene expression is stochastic [38], leading to significant cell-to-cell variation in protein numbers, which we model with a gamma distribution (Fig. S14B) [34, 35]. Therefore, the strong asymmetry of the fitness landscape predicts protein overabundance.

### D. Results: The RLTO model predicts overabundance is optimal for low-expression proteins

The optimal regulatory program for transcription and translation (*µ*_*m*_ and *ε* values) can be predicted analytically. The values are determined by a single global parameter, the relative load Λ, and the gene-specific threshold number *n*_*p*_. The threshold number is not directly observable experimentally; instead we predict the optimal overabundance *o*, defined as the ratio of the mean protein number to the threshold number:

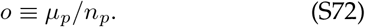

As shown in [11], by taking partial derivatives of the relative growth rate (Eq. S66) with respect to message number and translation efficiency, respectively, we can define the optimal overabundance:

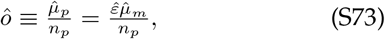

in the large multiplicity limit where the overall metabolic load is much smaller than the metabolic load for a single gene: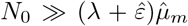. The optimal over-abundance can be rewritten to find the optimization condition for message number:

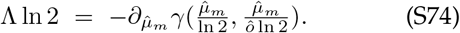

As seen in Fig. S14C, the RLTO model generically predicts that for a range of relative loads, the optimal protein fraction is overabundant (*o >* 1); however, overabundance is not uniform for all proteins, but rather depends on transcription. For highly-transcribed genes (*µ*_*m*_ ≫1), the overabundance is predicted to be quite small (*o* ≈ 1); however, for lowly-transcribed genes (message numbers approaching unity), the overabundance is predicted to be extremely high (*o* ≫ 1).

**FIG S14.**
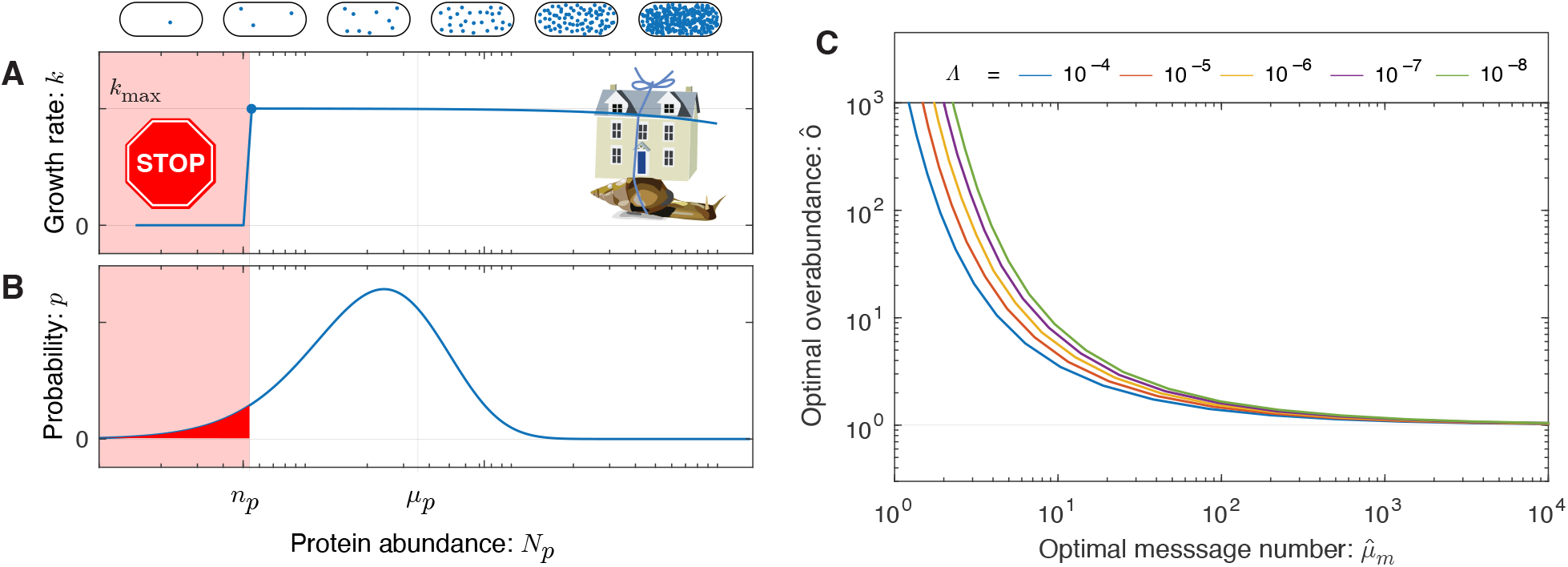
Panel A: The fitness landscape is asymmetric in the RLTO model. Motivated by single-cell growth data, cell fitness is modeled using the Robustness-Load Trade-Off model (RLTO). In the model, there is a metabolic cost of protein expression which favors low expression; however, growth arrests for protein number *N*_*p*_ smaller than the threshold level *n*_*p*_ (red). The relative metabolic cost of overabundance is small relative to the cost of growth arrest due to the large number of proteins synthesized, resulting in a highly asymmetric fitness landscape [11]. **Panel B: The gene expression process is stochastic**. There is significant cell-to-cell variation in protein abundance (*N*_*p*_) around the mean level (*µ*_*p*_). Even for mean expression levels significantly above the threshold level *n*_*p*_, some cells fall below threshold (red). The distribution in protein number is modeled using a gamma distribution [48]. **Panel C: Overabundance is predicted to optimize cell fitness**. The asymmetry of the fitness landscape drives the optimal protein expression level to be overabundant (*µ*_*p*_ *> n*_*p*_). The RLTO model makes a quantitative prediction of the optimal overabundance 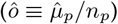 as a function of the message number *µ*_*m*_ and a global parameter, the relative load Λ ≈10^−5^ (red curve). Overabundance is predicted to be extremely high (*o* ≫ 1) for low expression genes (*µ*_*m*_ ≈1) and much closer to sufficiency (*o* ≈ 1) for high expression genes (*µ*_*m*_ ≫10). Although the optimal overabundance depends on the relative load Λ, its qualitative dependence is unchanged over orders of magnitude in variation of the parameter

### E. Discussion: Alternative hypothesis–Expression-dependent mutation rate

Could a gene-expression-dependent mutation rate explain the observed expression-dependence of overabundance? For instance, if the mutation rate were higher for low-expression genes, could this explain the observed increase in overabundance for low-expression genes? There are several strong arguments against this speculative alternative hypothesis. The expression dependence of the mutation rate has been studied in some detail already [54–57]. There is conflicting evidence in this previous work with some evidence indicating that mutation rates may rise with gene-expression [54, 56] and other work suggesting they decrease [55]. Furthermore, even if the mutation rates were to increase with mutation rate, this would not imply that the frequency of mutations at these loci would be higher. Due to the increased rate of mutation at these loci, more mutants would be created; however, their lifetime would be lower due to the same elevated mutation rate. These two competing effects are predicted to cancel, leading to a probability of mutants that is dependent only on the relative fitness of the mutants and the effective population size.

## 8. METHODS: STATISTICAL PROCEDURES

In this section, we provide a summary of statistical approaches that are common to the analyses in the paper.

### A. Maximum Likelihood Estimation

The maximum likelihood (*i*.*e*. minimum information) estimates (MLE) of the parameters are defined:

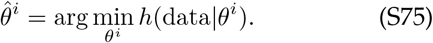

In all instances, these optimizations are performed numerically, either by direct minimization of the Shannon information (*h*), or for normal models, by least-squares minimization.

### B. Parametric uncertainty estimates

To estimate the parameter uncertainty in the analysis of datasets, we use the Cramer-Rao bound to estimate of the uncertainty from the Fisher information [28]:

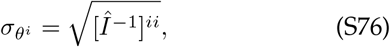

where 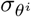 is the estimate of the standard error for parameter *θ*^*i*^, 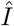 is the estimator of the Fisher information, and 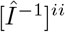 is the *ii* component of the inverse Fisher information matrix. For each statistical model, we describe how the Fisher information is estimated in detail (Hessian or Jacobian *etc*).

### C. Null-hypothesis-testing approach

For null-hypothesis testing, we define two sequential null-hypothesis tests of nested statistical models. If the initial null hypothesis is rejected, we then interpret the initial alternative hypothesis as the updated null hypothesis and adopt the remaining model as the alternative hypothesis. For each test, we will use a Likelihood Ratio Test (LRT) where we define the test statistic *λ* in terms of the Shannon information:

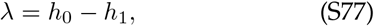

where *h*_0_ and *h*_1_ are the Shannon information for the null and alternative hypotheses respectively. We will assume the Wilks’ theorem: *I*.*e*. the test statistic Λ under the null hypothesis will have a chi-squared distribution [58, 59]:

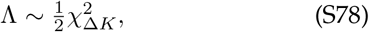

where the degrees-of-freedom Δ*K* = 1 is equal to the difference between the dimension of the alternative and null models. (The factor of 1/2 appears in this equation, since the test statistic is defined by the Shannon information difference rather than the deviance [28].) The p-value can then be computed:

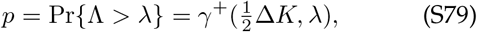

where *γ*^+^ is the upper regularized incomplete gamma function (Eq. S95), Δ*K* = 1 is the difference in model dimensions, and *λ* is the test statistic [28].

## 9. DISTRIBUTIONS AND CONVENTIONS

### A. Gamma distribution conventions

There are a number of conflicting conventions for the gamma function and distribution arguments. We will use those defined on Wikipedia and the CRC Encyclopedia of Mathematics [60].

The gamma distributed random variable *X* will be written:

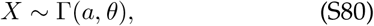

where *a* is the shape parameter and *θ* is the scale parameter. The PDF of the distribution is:

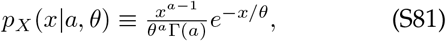

where Γ(*a*) is the gamma function. The CDF is therefore:

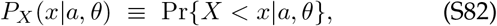

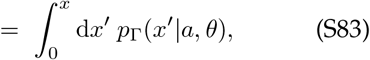

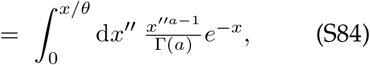

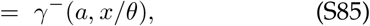

where *γ*^−^ is the regularized lower incomplete gamma function. The survival function is:

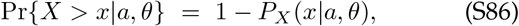

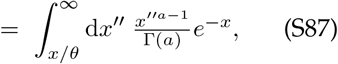

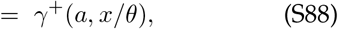

where *γ*^+^ is the regularized upper incomplete gamma function.

### B. Chi-squared distribution conventions

In statistical null hypothesis testing, the chi-squared distribution arises in the context of the Likelihood Ratio Test (LRT). Let *Y* be distributed like a chi-squared with *k* degrees of freedom:

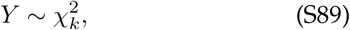

where the PDF is:

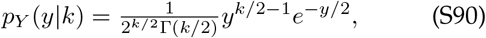

where Γ is the gamma function. The CDF is therefore:

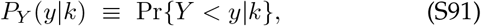

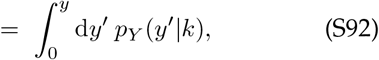

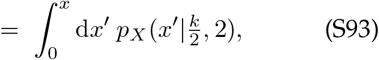

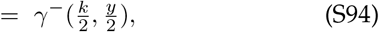

where *p*_*X*_ is the PDF of the gamma distribution (Eq. S81) and *γ*^−^ is the regularized lower incomplete gamma function. The survival function is:

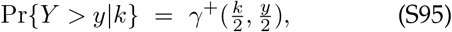

where *γ*^+^ is the regularized upper incomplete gamma function.

## 10. DESCRIPTION OF SUPPLEMENTARY DATA

### A. Data Tables

**Data S1**: Overabundance for all genes as measured by TFNseq analysis. The original TFNseq data was previously generated by the Manoil lab [18]. Format: Open Document Format (xlsx).

**Data S2**: A list of essential genes ranked by overabundance. Format: Open Document Format (xlsx).

**Data S3**: Representative single-cell imaging-based cell cytometry data for wild-type *A. baylyi* proliferating on minimal media (Km^-^) from a single progenitor cell (Sec. 1 D). Format: Open Document Format (xlsx).

**Data S4**: Representative single-cell imaging-based cell cytometry data for *A. baylyi* Δ*IS* proliferating on minimal media (Km^+^) from a single progenitor cell in a knockout-depletion experiment (Sec. 1 D). Format: Open Document Format (xlsx).

**Data S5**: Representative single-cell imaging-based cell cytometry data for *A. baylyi* Δ*dnaA* proliferating on minimal media (Km^+^) from a single progenitor cell in a knockout-depletion experiment (Sec. 1 E). Format: Open Document Format (xlsx).

**Data S6**: Representative single-cell imaging-based cell cytometry data for *A. baylyi* Δ*dnaN* proliferating on minimal media (Km^+^) from a single progenitor cell in a knockout-depletion experiment (Sec. 1 F). Format: Open Document Format (xlsx).

**Data S7**: Representative single-cell imaging-based cell cytometry data for *A. baylyi* Δ*murA* proliferating on minimal media (Km^+^) from a single progenitor cell in a knockout-depletion experiment (Sec. 1 H). Format: Open Document Format (xlsx).

**Data S8**: Representative single-cell imaging-based cell cytometry data for *A. baylyi* Δ*ftsN* proliferating on minimal media (Km^+^) from a single progenitor cell in a knockout-depletion experiment (Sec. 1 G). Format: Open Document Format (xlsx).

### B. Annotated sequences

**Data S9**: The annotated sequence of the DnaN fluorescent fusion *YPet-dnaN*. Format: Genbank file.

### C. Supplemental movies

**Movie S1**: Wild-type *A. baylyi* proliferating on minimal media (Km^-^). Frame rate: 1 frame/2 min. (Sec. 1 D.) Raw images.

**Movie S2**: Wild-type *A. baylyi* proliferating on minimal media (Km^-^). Frame rate: 1 frame/2 min. (Sec. 1 D.) Annotated/segmented images.

**Movie S3**: *A. baylyi* Δ*IS* proliferating on minimal media (Km^+^) in a knockout-depletion experiment. Frame rate: 1 frame/3 min. (Sec. 1 D.) Raw images.

**Movie S4**: *A. baylyi* Δ*IS* proliferating on minimal media (Km^+^) in a knockout-depletion experiment. Frame rate: 1 frame/3 min. (Sec. 1 D.) Annotated/segmented images.

**Movie S5**: *A. baylyi* Δ*dnaA* proliferating on minimal media (Km^+^) in a knockout-depletion experiment. Frame rate: 1 frame/2 min. (Sec. 1 E.) Raw images.

**Movie S6**: *A. baylyi* Δ*dnaA* proliferating on minimal media (Km^+^) in a knockout-depletion experiment. Frame rate: 1 frame/2 min. (Sec. 1 E.) Annotated/segmented images.

**Movie S7**: *A. baylyi* Δ*dnaN* proliferating on minimal media (Km^+^) in a knockout-depletion experiment. Frame rate: 1 frame/9 min. (Sec. 1 F.) Raw images.

**Movie S8**: *A. baylyi* Δ*dnaN* proliferating on minimal media (Km^+^) in a knockout-depletion experiment. Frame rate: 1 frame/9 min. (Sec. 1 F.) Annotated/segmented images.

**Movie S9**: *A. baylyi* Δ*murA* proliferating on minimal media (Km^+^) in a knockout-depletion experiment. Frame rate: 1 frame/2 min. (Sec. 1 H.) Raw images.

**Movie S10**: *A. baylyi* Δ*murA* proliferating on minimal media (Km^+^) in a knockout-depletion experiment. Frame rate: 1 frame/2 min. (Sec. 1 H.) Annotated/segmented images.

**Movie S11**: *A. baylyi* Δ*ftsN* proliferating on minimal media (Km^+^) in a knockout-depletion experiment. Frame rate: 1 frame/2 min. (Sec. 1 G.) Raw images.

**Movie S12**: *A. baylyi* Δ*ftsN* proliferating on minimal media (Km^+^) in a knockout-depletion experiment. Frame rate: 1 frame/2 min. (Sec. 1 G.) Annotated/segmented images.

## References

[1] Alberts B, et al. (2002) Molecular Biology of the Cell. (Garland), 4th edition.

[2] Dekel E, Alon U (2005) Optimality and evolutionary tuning of the expression level of a protein. Nature 436(7050):588–92.

[3] Lalanne JB, Parker DJ, Li GW (2021) Spurious regulatory connections dictate the expression-fitness landscape of translation factors. Mol Syst Biol 17(4):e10302.

[4] Lengeler JW, Drews G, Schlegel HG, eds. (1998) Biology of the Prokaryotes. (Georg Thieme Verlag, Rüdigerstrasse 14, D–70469 Stuttgart, Germany).

[5] Kafri M, Metzl-Raz E, Jonas F, Barkai N (2016) Rethinking cell growth models. FEMS Yeast Res 16(7).

[6] Hausser J, Mayo A, Keren L, Alon U (2019) Central dogma rates and the trade-off between precision and economy in gene expression. Nat Commun 10(1):68.

[7] Belliveau NM, et al. (2021) Fundamental limits on the rate of bacterial growth and their influence on proteomic composition. Cell Syst 12(9):924–944.e2.

[8] Peters JM, et al. (2016) A comprehensive, CRISPR-based functional analysis of essential genes in bacteria. Cell 165(6):1493–1506.

[9] Donati S, et al. (2021) Multi-omics analysis of CRISPRi-knockdowns identifies mechanisms that buffer decreases of enzymes in E. coli metabolism. Cell Syst 12(1):56–67.e6.

[10] Baba T, Huan HC, Datsenko K, Wanner BL, Mori H (2008) The applications of systematic in-frame, singlegene knockout mutant collection of Escherichia coli K-12. Methods Mol Biol 416:183–94.

[11] Lo TW, Choi HJ, Huang D, Wiggins PA (2024) Noise robustness and metabolic load determine the principles of central dogma regulation. Science Advances 10(34).

[12] McGinness KE, Baker TA, Sauer RT (2006) Engineering controllable protein degradation. Mol Cell 22(5):701–7.

[13] Davis JH, Baker TA, Sauer RT (2011) Small-molecule control of protein degradation using split adaptors. ACS Chem Biol 6(11):1205–13.

[14] Cameron DE, Collins JJ (2014) Tunable protein degradation in bacteria. Nat Biotechnol 32(12):1276–81.

[15] Liu X, et al. (2017) High-throughput CRISPRi phenotyping identifies new essential genes in Streptococcus pneumoniae. Mol Syst Biol 13(5):931.

[16] Franks SNJ, Heon-Roberts R, Ryan BJ (2024) CRISPRi: a way to integrate iPSC-derived neuronal models. Biochem Soc Trans 52(2):539–551.

[17] Bailey J, et al. (2019) Essential gene deletions producing gigantic bacteria. PLoS Genet 15(6):e1008195.

[18] Gallagher LA, Bailey J, Manoil C (2020) Ranking essential bacterial processes by speed of mutant death. Proc Natl Acad Sci U S A 117(30):18010–18017.

[19] Mangiameli SM, Merrikh CN, Wiggins PA, Merrikh H (2017) Transcription leads to pervasive replisome instability in bacteria. Elife 6.

[20] Mangiameli SM, Veit BT, Merrikh H, Wiggins PA (2017) The replisomes remain spatially proximal throughout the cell cycle in bacteria. PLoS Genet 13(1):e1006582.

[21] Mangiameli SM, Cass JA, Merrikh H, Wiggins PA (2018) The bacterial replisome has factory-like localization. Curr Genet 64(5):1029–1036.

[22] Kuwada NJ, Traxler B, Wiggins PA (2015) Genome-scale quantitative characterization of bacterial protein localization dynamics throughout the cell cycle. Mol Microbiol 95(1):64–79.

[23] Cooper GM (2000) The Cell: A Molecular Approach. 2nd edition. (Sinauer Associates 2000).

[24] Lemon KP, Grossman AD (1998) Localization of bacterial dna polymerase: evidence for a factory model of replication. Science 282(5393):1516–9.

[25] Reyes-Lamothe R, Sherratt DJ, Leake MC (2010) Stoichiometry and architecture of active DNA replication machinery in Escherichia coli. Science 328(5977):498–501.

[26] Cutler KJ, et al. (2022) Omnipose: a high-precision morphology-independent solution for bacterial cell segmentation. Nat Methods 19(11):1438–1448.

[27] de Berardinis V, et al. (2008) A complete collection of single-gene deletion mutants of acinetobacter baylyi adp1. Mol Syst Biol 4:174.

[28] Cox DR, Hinkley DV (1974) Theoretical Statistics. (Chapman & Hall, London, England).

[29] Autret S, Levine A, Holland IB, Séror SJ (1997) Cell cycle checkpoints in bacteria. Biochimie 79(9-10):549–54.

[30] Hawkins JS, et al. (2020) Mismatch-CRISPRi reveals the co-varying expression-fitness relationships of essential genes in Escherichia coli and Bacillus subtilis. Cell Syst 11(5):523–535.e9.

[31] Keren L, et al. (2016) Massively parallel interrogation of the effects of gene expression levels on fitness. Cell 166(5):1282–1294.e18.

[32] Bosch B, et al. (2021) Genome-wide gene expression tuning reveals diverse vulnerabilities of M. tuberculosis. Cell 184(17):4579–4592.e24.

[33] Nelson DL, Cox MM (2017) Lehninger principles of biochemistry. (W.H. Freeman, New York, NY), 7 edition.

[34] Paulsson J, Ehrenberg M (2000) Random signal fluctuations can reduce random fluctuations in regulated components of chemical regulatory networks. Phys Rev Lett 84(23):5447–50.

[35] Friedman N, Cai L, Xie XS (2006) Linking stochastic dynamics to population distribution: an analytical framework of gene expression. Phys Rev Lett 97(16):168302.

[36] Lambert G, Kussell E (2014) Memory and fitness optimization of bacteria under fluctuating environments. PLoS Genet 10(9):e1004556.

[37] Mori M, Schink S, Erickson DW, Gerland U, Hwa T (2017) Quantifying the benefit of a proteome reserve in fluctuating environments. Nat Commun 8(1):1225.

[38] Raser JM, O’Shea EK (2005) Noise in gene expression: origins, consequences, and control. Science 309(5743):2010–3.

[39] Weichart D, Querfurth N, Dreger M, Hengge-Aronis R (2003) Global role for ClpP-containing proteases in stationary-phase adaptation of Escherichia coli. J Bacteriol 185(1):115–25.

[40] Goldberg AL, St John AC (1976) Intracellular protein degradation in mammalian and bacterial cells: Part 2. Annu Rev Biochem 45:747–803.

[41] Barbe V, et al. (2004) Unique features revealed by the genome sequence of Acinetobacter sp. ADP1, a versatile and naturally transformation competent bacterium. Nucleic Acids Res 32(19):5766–79.

[42] Miller J (1972) Experiments in Molecular Genetics, Bacterial genetics E. coli. (Cold Spring Harbor Laboratory).

[43] Chang AC, Cohen SN (1978) Construction and characterization of amplifiable multicopy DNA cloning vehicles derived from the P15A cryptic miniplasmid. J Bacteriol 134(3):1141–56.

[44] Lo TW, et al. (2024) OmniSegger GitHub.

[45] Burnham KP, Anderson DR (2004) Multimodel inference: Understanding AIC and BIC in model selection. Socialogical Methods & Research 33(2):261–304.

[46] Karp PD, et al. (2023) The EcoCyc database (2023). EcoSal Plus 11(1):eesp–0002–2023.

[47] Culviner PH, Guegler CK, Laub MT (2020) A simple, costeffective, and robust method for rRNA depletion in RNA-Sequencing studies. mBio 11(2).

[48] Taniguchi Y, et al. (2010) Quantifying E. coli proteome and transcriptome with single-molecule sensitivity in single cells. Science 329(5991):533–8.

[49] Crick F (1970) Central dogma of molecular biology. Nature 227(5258):561–3.

[50] Hargrove JL, Schmidt FH (1989) The role of mRNA and protein stability in gene expression. FASEB J 3(12):2360– 70.

[51] Koch AL, Levy HR (1955) Protein turnover in growing cultures of Escherichia coli. J Biol Chem 217(2):947–57.

[52] Martin-Perez M, Villén J (2017) Determinants and regulation of protein turnover in yeast. Cell Syst 5(3):283–294.e5.

[53] Scott M, Gunderson CW, Mateescu EM, Zhang Z, Hwa T (2010) Interdependence of cell growth and gene expression: origins and consequences. Science 330(6007):1099– 102.

[54] Merrikh H, Zhang Y, Grossman AD, Wang JD (2012) Replication-transcription conflicts in bacteria. Nat Rev Microbiol 10(7):449–58.

[55] Martincorena I, Seshasayee ASN, Luscombe NM (2012) Evidence of non-random mutation rates suggests an evolutionary risk management strategy. Nature 485(7396):95– 8.

[56] Chen X, Zhang J (2013) No gene-specific optimization of mutation rate in escherichia coli. Mol Biol Evol 30(7):1559– 62.

[57] Jee J, et al. (2016) Rates and mechanisms of bacterial mutagenesis from maximum-depth sequencing. Nature 534(7609):693–6.

[58] Wilks SS (1938) The large-sample distribution of the likelihood ratio for testing composite hypotheses. The Annals of Mathematical Statistics. 9(1):60–62.

[59] Casella G, Berger R (2001) Statistical Inference. (Duxbury Resource Center).

[60] Weisstein EW (2009) CRC Encyclopedia of Mathematics. (Chapman & Hall/CRC).

